# Schistosomiasis parasite enhance transmission rates via interfacial swimming

**DOI:** 10.1101/2025.05.13.653683

**Authors:** Melanie Hannebelle, Ian Ho, Alassane N’diaye, Manu Prakash

## Abstract

Schistosomiasis is a disease of poverty that affects over 240 million people worldwide despite an ecological paradox: cercariae larvae are short-lived (12 hours), sparse in the water body, and entrained by flows exceeding their swimming speeds - conditions that should limit host-finding. We resolve this paradox by demonstrating that cercariae utilize a physics-based strategy: they actively accumulate at the air-water interface, transforming inefficient three-dimensional search into effective two-dimensional exploration, increasing host-encounter efficiency by a thousand fold. Our biophysical models demonstrate that this behavior emerges from weight-asymmetric morphology, enabling an embodied algorithm for surface-seeking swimming modes without complex neural control. Surface-swimming cercariae benefit from near-zero vertical flows near the surface, allowing them to remain there while exploiting wind-driven currents for long-distance dispersal. We identify the air-water interface as a critical transmission micro-habitat, suggesting new physical intervention strategies targeting this interface to mitigate transmission of this highly infectious disease.

---

Schistosomiasis is a neglected tropical disease affecting approximately 240 million people worldwide, predominantly low-income populations of sub-Saharan Africa in communities living near freshwater ecosystems (*1*) (movie S1). The parasite infects humans by entering human skin via contact and causes chronic inflammation and organ damage, contributing to anemia, malnutrition, impaired cognitive development, and increased vulnerability to other infections (*2, 3*).

Despite the mass drug administration of praziquantel treatment, reinfection rates exceed 40% annually in some endemic areas (*4–6*) (fig. S1). Alternative control approaches face significant challenges: molluscicides pose environmental risks (*7*), biological control methods present logistical difficulties (*8–10*), and water infrastructure projects often expand the parasite habitats (*11–13*). A recent outbreak in southern Europe highlights risks exacerbated by climate change and human migration (*14*).

The transmission cycle of schistosomes begins when parasite eggs enter freshwater via human excreta. Schistosomes release thousands of eggs daily through human urine (*Schistosoma haematobium*) or stool (*Schistosoma mansoni* and *Schistosoma japonicum*) (*15*). These hatch into miracidia, which infect specific species of freshwater snails (fig.1A). Inside snails, parasites multiply asexually, releasing thousands of free-swimming cercariae (*16*) per day that seek human hosts in an open water body such as a lake, river or pond, penetrate skin, and complete their lifecycle by maturing and producing new eggs (fig. 1A) inside a human host.

**Figure 1:**
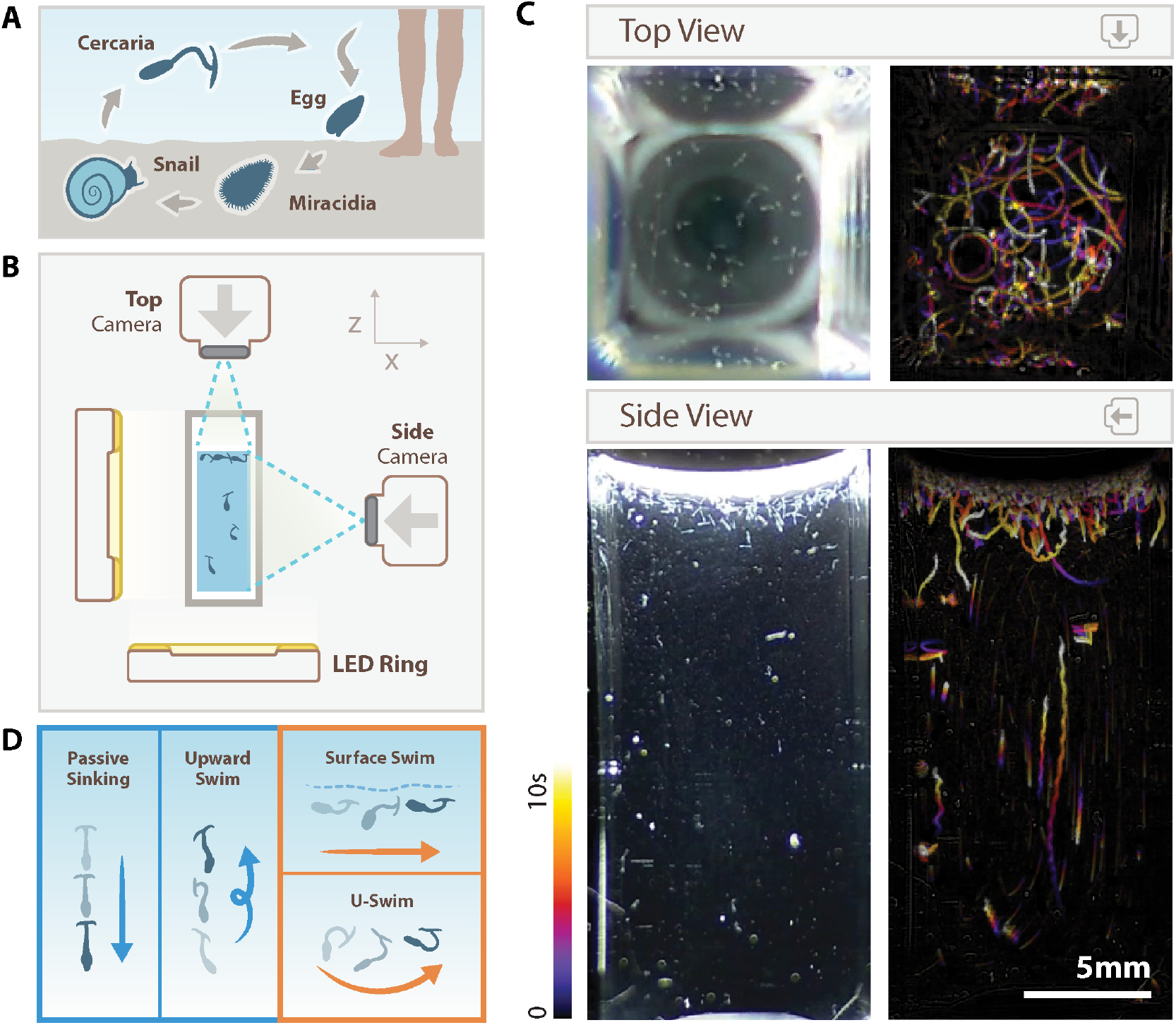
Schistosome cercariae adopt a discrete set of swimming modes. **A** Schematic of the life cycle of schistosoma parasites in the water stages. **B** Experimental imaging setup with two cameras, imaging a cuvette with cercariae from the top and from the side. **C** Bright field image (left) and temporal color-coded projection of the displacement of cercariae (right) within a 1 cm wide square cuvette. **D** Schematic representation of the swimming trajectories observed experimentally. Head-first swimming is not represented here because it is associated with short distance displacement near a potential host.

Surprisingly, field studies reveal remarkably low cercarial densities in natural environments, measured to be only 0.0016-0.032 cercariae per liter in sampled sites in Kenya, with a maximum of 0.5 cercariae per liter (*17,18*). In addition, cercariae show no directed motion toward the skin compounds, indicating absence of chemotaxis (*19–21*). Instead, they exhibit chemokinesis: modification of swimming patterns in response to a stimulus (*19–21*). Yet they successfully infect hundreds of millions of people. Therefore, despite decades of research into schistosomiasis biology and control, a fundamental yet underappreciated ecological puzzle remains: how do cercariae successfully infect human hosts given such dilute concentrations and lack of target-oriented navigation?

We address this paradox through integrated multi-scale imaging both in lab and field observations, and physical hydrodynamic models(fig. S2). We demonstrate that cercariae exploit a previously unrecognized strategy: active accumulation at the air-water interface that arises naturally from the link between the underlying parasite geometry and underlying hydrodynamics. This adaptation transforms a three-dimensional search problem into a two-dimensional surface exploration, enhancing host-encounter efficiency by orders of magnitude. Our findings identify the air-water interface as a critical transmission niche and suggest new physical intervention strategies to complement existing control methods for this disease that still remains unchecked.

## The paradox of cercarial transmission

Cercariae must navigate from infected snails to humans within their 10-20 hour lifespan (*22*). This window narrows as humans occupy water bodies briefly, limiting transmission opportunity to minutes or hours. A simple analysis highlights this challenge: cercariae swim at approximately 1 mm/s (*23*), explore with a cross-sectional area of roughly 1 mm^3^, and exist at field-measured peak concentrations of 0.5 cercariae per liter (*17*). Under these conditions, systematically exploring a cubic meter of water would require approximately 1m^3^/(500cerc./m^3^)/(1mm^3^/s) = 23 days, far exceeding their lifespan. This paradox demonstrates that volumetric exploration alone would make transmission virtually impossible, suggesting cercariae must employ a more efficient host-seeking strategy.

### Cercarial swimming trajectories form a distinctive locomotion repertoire

Imaging cercarial movement (fig. 1B,C; movie S2; fig. S4) reveals four distinct swimming patterns (fig. 1D). Our previous work documented alternating phases of active upward swimming and passive sinking (*23, 24*), which we observe as well, particularly in cuvettes less than 3mm thick (fig. S3). When cercariae are given the space to swim in 3 dimensions, we identify two additional, previously uncharacterized modes: surface swimming (horizontal movement at the air-water interface) and U-swimming (parabolic-like trajectories away from the surface) (fig.1D). A fifth mode, headfirst swimming, occurs only within approximately 1 mm of potential hosts (*23*) and represents the terminal penetration phase rather than the search component, so we exclude it from our current analysis of host-seeking behavior.

### A biophysical model captures cercarial swimming modes

We sought to understand why cercarial locomotion is constrained to four distinct patterns and determine whether they confer an advantage in the context of host-search. Our biophysical analysis reveals that these diverse trajectories emerge from physical constraints associated with the weight asymmetry between the cercarial head and tail.

We first develop a reduced order biophysical model of cercaria under the influence of gravity (see Supplementary Text), representing each cercaria as a rigid elongated object with weight asymmetry (fig. 2A) (*23*). The weight generates a downward force *F*_*g*_ and an asymmetry-based torque *T*_*g*_ = *T*_*g*0_ · cos(*θ*) that reorients the cercaria vertically. Swimming is simplified to a constant self-propelling force *F*_*p*_, while the air-water interface interaction is modeled as a downward elastic force arising from interfacial deformation *F*_*s*_ generating a torque *T*_*s*_ (fig. 2B). This simple model in low Reynolds number hydrodynamics predicts trajectories under the influence of gravity captured by the following equations of motion,

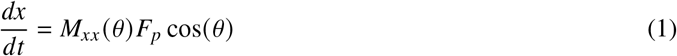

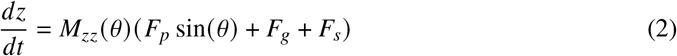

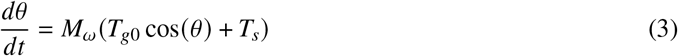

where mobility coefficients *M*_*xx*_, *M*_*zz*_, and *M*_*ω*_ derive from cercaria geometry and fluid properties (Supplementary Text).

**Figure 2:**
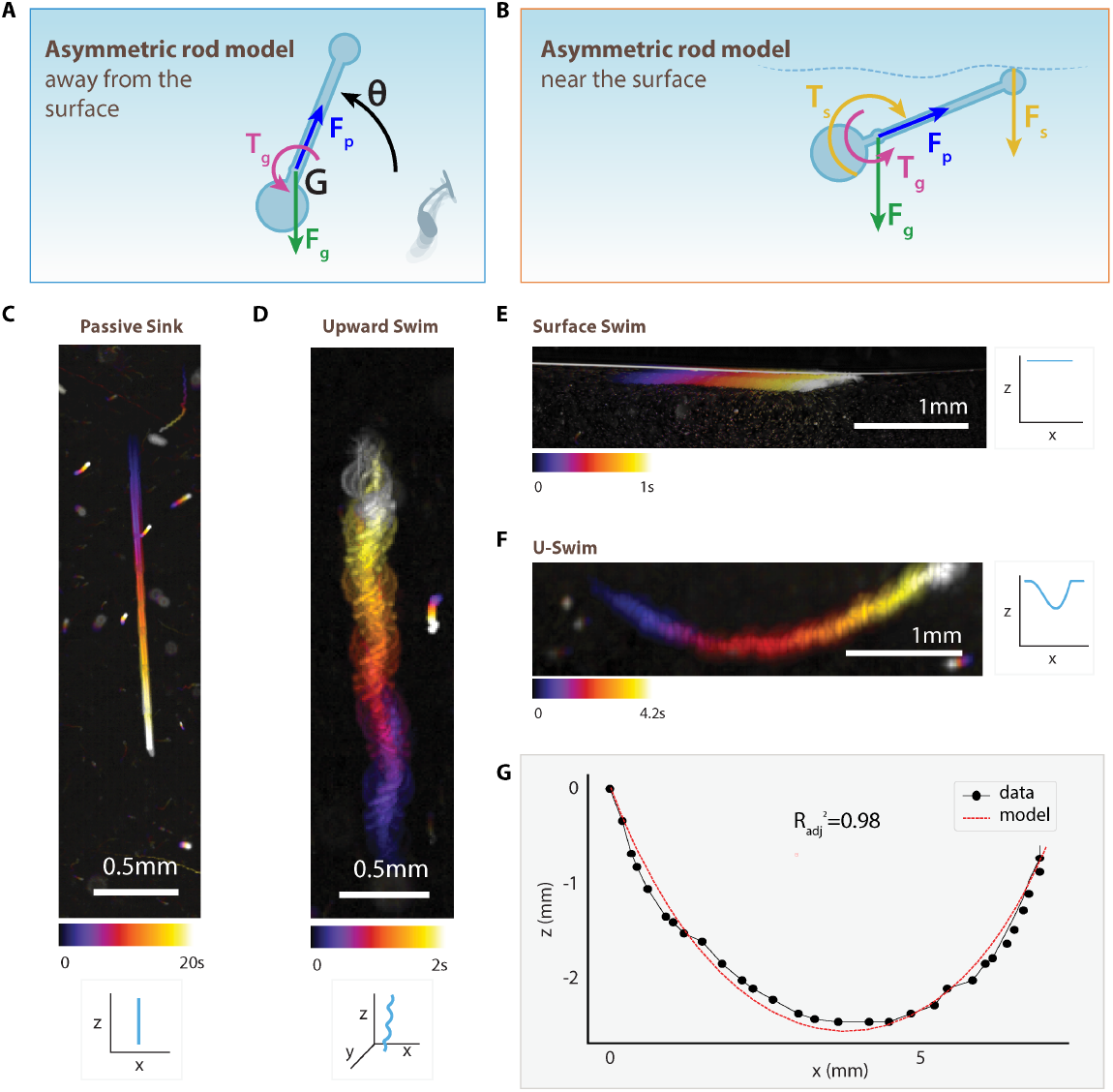
An asymmetric rod model reproduces the four swimming modes of cercariae. **A** Schematic representation of the asymmetric rod model, for a cercaria away from the surface. *F*_*g*_ is the gravitational force, *T*_*g*_ the torque associated with the weight asymmetry between the head and the tail of the cercaria, *F*_*p*_ a constant force recapitulating the swimming gait. **B** Model of a cercaria at the air-water interface. *F*_*s*_ and *T*_*s*_ represent respectively the elastic force and the torque associated with the surface interaction. **C** Simulation with *F*_*p*_ = 0 and experimental trajectory of a sinking cercaria. **D** Simulation away from the surface and experimental trajectory of an upward swimming cercaria. The spiral motion is obtained in 3D when *F*_*p*_ is slightly off axis relative to the axis of the swimmer in the model. **E** Simulation at the water surface and experimental trajectory of a surface swimming cercaria. **F** Simulation with *θ* (*t* = 0) ∈]0, −*π*/2[and experimental trajectory of a u-swimming cercaria. **G** Representative u-swim experimental curve (black) and fit with the asymmetric rod model (red). The adjusted coefficient of determination 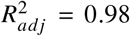 means that 98% of the data variability is explained by the variation of the inputs of the model.

Remarkably, this simple physical model reproduces all four swimming modes without requiring neural control mechanisms. These correspond to distinct solutions: (1) passive sinking when *F*_*p*_ = 0 (fig. 2D); (2) upward swimming when starting vertically (*θ* (*t* = 0) = *π*/2) (fig. 2D); (3) surface swimming when maintaining surface contact with *F*_*s*_ ≠ 0 (fig. 2E); and (4) U-swimming when starting non-vertically (*θ* (*t* = 0) ≠ *π*/2) away from the surface (fig. 2F). The simulation results (fig. 2G, fig. S8, fig. S10) show excellent agreement with experimental observations with an adjusted *R*^2^ = 0.96 ± 0.04 for *N* = 14 fitted curves (fig. S9), which validates our model.

The model shows trajectory depends solely on initial position and orientation (fig. S11A,B). A critical finding is that the maximum lateral displacement of cercariae during U-swimming is constrained to 9.3mm, occurring when the cercaria begins in an inverted position (fig. S11C). This constraint exists because lateral movement stops once gravitational torque reorients cercariae vertically. Our volumetric exploration estimate (23 days) thus underestimates the actual challenge, as the lateral exploration is physically limited to about one centimeter maximum while away from the water surface.

### Buoyant torque enables aggregation towards the air-water interface for cercariae

While our asymmetric rod model successfully reproduces the four swimming modes, the elastic force *F*_*s*_ and torque *T*_*s*_ poorly approximate the complex hydrodynamic interactions with the air-water interface. To explore these interactions in detail, we developed a two-dimensional dumbbell model incorporating single-sphere hydrodynamic interactions with an interface (fig. 3A) (*26–33*). This approach uses time-varying sphere radii and rod length to achieve self-propulsion, accurately representing the non-reciprocal deformations that enable swimming at low Reynolds number (fig. 3B) (*23, 34–37*). Although slender–body formulations exist for swimmers near interfaces, they are either restricted to specific orientations and moderate separations where lubrication stresses are weak (*38–40*), with no uniformly valid theory that spans arbitrary orientations and gap sizes (*41,42*). We therefore adopt a coarse-grained shape-space model (*34*) that isolates the leading-order impacts of buoyant torque and a planar, impenetrable interface while sidestepping the unresolved near-field complexities. Specifically,

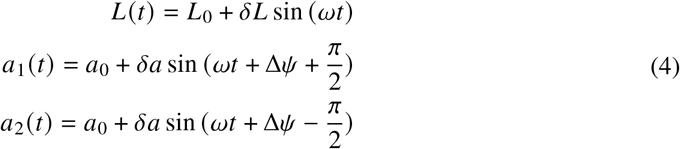

where *L*_0_ and *a*_0_ represent the mean length and radius, respectively; *δL* and *δa* denote the corresponding amplitudes; *ω* is the angular beating frequency; and Δ*ψ* is a phase shift that modulates the instantaneous velocity of the dumbbell. More details of the model can be found in the Supplementary Text. Crucially, we find that in the absence of body forces or torques, modeled cercariae experience fundamentally different hydrodynamic interactions depending on the interface type: they are attracted towards an air-water interface but repelled from a solid-water boundary (fig. 3C). Our work builds upon the vast amount of literature describing hydrodynamic attraction, repulsion, and reorientation towards/away from surfaces (*38, 43–60*); here, we find a new consequence of interfacial swimming for disease dispersal.

**Figure 3:**
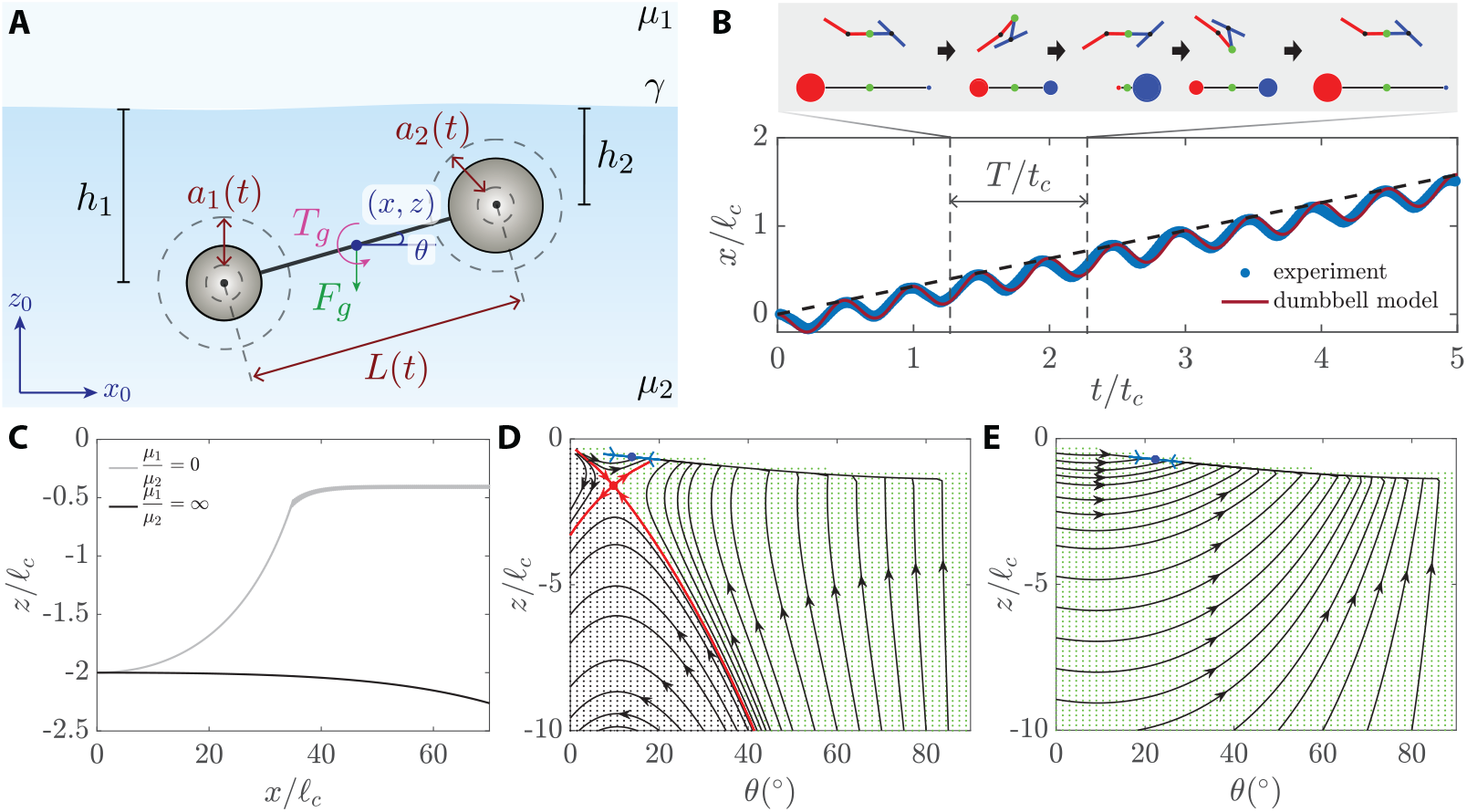
A dumbbell model reveals hydrodynamic attraction to the air-water interface. **A**. Our dumbbell model represents cercariae as two spheres with time-varying radii *a*_1,2_(*t*) connected by a drag-less rod with time-varying length *L* (*t*). The dumbbell swims in a fluid of viscosity *µ*_2_ near an interface with a fluid of viscosity *µ*_1_, with surface tension *γ*. A gravitational torque *T*_*g*_ and force *F*_*g*_ act at the geometric center. **B** Time-varying sphere radii and rod length reproduce the periodic, non-reciprocal deformations characteristic of the *T-swimmer* gait (*23*), enabling propulsion at low Reynolds number. Model parameters were calibrated using experimental data from horizontally swimming cercariae. **C** In the absence of body force (*T*_*g*0_ = *F*_*g*_ = 0), a swimmer initially parallel to the interface experiences fundamentally different behaviors depending on the interface type: attraction toward an air-water interface (*µ*_1_/*µ*_2_ ≈ 0) but repulsion from a solid-water interface (*µ*_1_/*µ*_2_ *→* ∞). **D-E** Phase portraits of the Poincaré map in *z*-*θ* space (*z* ∈ [0, −10] and *θ* ∈ [0, 90^°^)) reveal linearly stable fixed points (blue markers) which corresponds to orbitally stable limit cycles in the full dynamics. **D** The absence of buoyant torque (*T*_*g*0_ = 0) produces an additional saddle point whose separatrix (red) diverts part of phase space. **E** Introducing a physiologically estimated buoyant torque (*T*_*g*0_ > 0) removes the saddle, so all trajectories converge to the fixed point. All torques are reported in nondimensional units. Background shading shows Monte-Carlo sampling of initial conditions (*25*): green points converge to the fixed point, grey do not.

We next focus on cercariae initialized near the air–water interface and ask whether buoyant torque (*T*_*g*0_) allows them to migrate towards the surface. To address this, we sweep the phase space of the Poincaré map (fig. 3D–E) within a bounded domain and identify linearly stable fixed points to which swimming trajectories converge to a steady height and orientation relative to the interface when averaged over one beat cycle. Importantly, the size of the basin of attraction depends on the buoyant torque. For *T*_*g*0_ = 0, an additional saddle point appears whose separatrix diverts part of phase space away from the stable node (fig. 3D). Introducing a realistic buoyant torque *T*_*g*0_ > 0 eliminates the saddle (fig. 3E), and every trajectory launched within the range of physiologically relevant initial conditions converges to the fixed point. Thus, the coupled action of buoyant torque, an impenetrable interface and near-interface hydrodynamics generates an attracting swimming state that robustly guides cercariae to the surface, regardless of initial position or orientation.

### Surface swimming is the most frequent swimming mode

This mathematical prediction of surface preference is strongly supported by our experimental observations of cercariae behavior in natural field conditions in Senegal: in a naturalistic 15×15×15cm aquarium with water, vegetation, and infected snails (*B. glabrata*) freshly collected in Mbakhana, Senegal, cercariae formed a concentrated layer at the water surface (fig. 4A, movie S3). Analysis in 1×1×4 cm cuvettes confirmed this surface preference, with 86 ± 3% of cercariae (3 biological replicates) positioned within 1 mm of the air-water interface (fig. S5A, movie S4) and tracking of a single cercaria showed a similar value of 89% of a 1000-second period in this zone (fig.S5B, movie S5).

**Figure 4:**
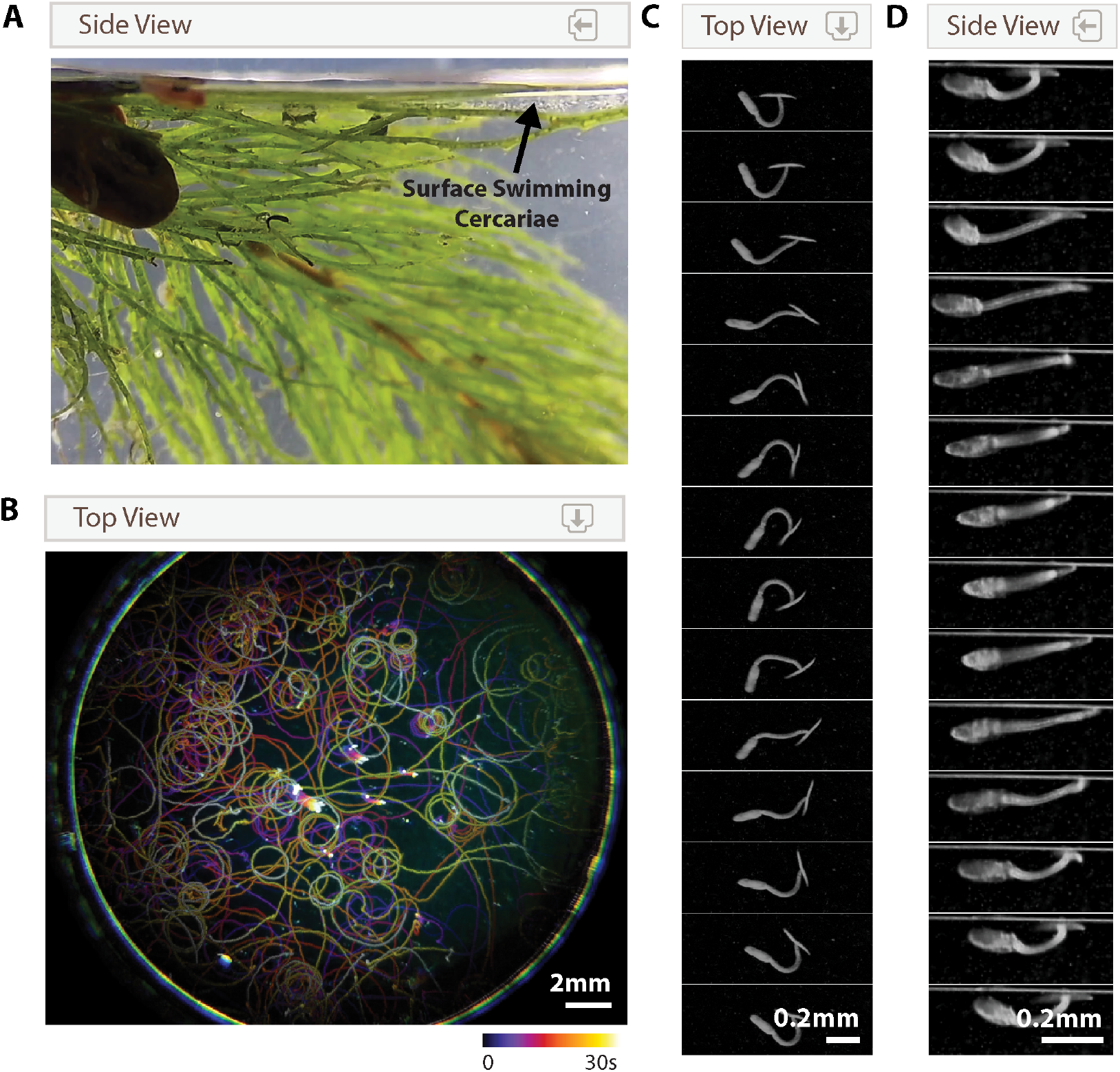
Surface swimming is the preferred swimming mode. **A** Picture taken from the side of an aquarium containing an infected snail, water and vegetation both sampled from Mbakhana, Senegal. Cercariae are visible as bright white spots predominantly near the water surface. **B** Temporal color-coded projection of the trajectories of cercariae, seen from above. The circular trajectories of the cercariae at the surface are visible. **C** Time-lapse high resolution imaging of a surface swimming cercaria, seen from above. Total duration: 70 ms. **D** Time-lapse high resolution imaging of a surface swimming cercaria, seen from the side. Total duration: 80ms.

Surface-swimming cercariae exhibit distinctive circular trajectories (fig. 4B, fig. S6A, movie S5, movie S6), predominantly anticlockwise (fig. S7A), with periodic sinking episodes (fig. S6B-D). These circles have a median radius of 4.4mm, with swimming speeds of 1.0 ± 0.2mm · s^−1^ and frequencies of 13.0 ± 0.6Hz (fig. S7B-E). Though their swimming gait resemble upward swimming (*23*), surface swimmers maintain an orientation nearly parallel to the interface (10.1±0.8^°^ inclination; fig. 4C,D, fig.S7C) without the axial spiral rotation typical of volumetric swimming, their tail tips appearing to maintain contact with the air-water interface (movie S7-S8).

### Cercariae are within water flows that exceed their swimming speed

A critical yet overlooked aspect of cercarial transmission is their navigation through complex flow environments. Our underwater imaging in natural conditions in Mbakhana, Senegal (fig. 5A, fig. S14, movie S9), revealed ambient flow velocities averaging 1.5mm/s in transmission habitats, exceeding the cercarial maximum swimming capacity of 0.7 mm/s (*23*) (fig. 5B). This velocity differential shows cercariae cannot maintain position against currents through locomotion alone, suggesting they must exploit specialized mechanisms to remain in host-accessible regions despite being outpaced by environmental flows.

**Figure 5:**
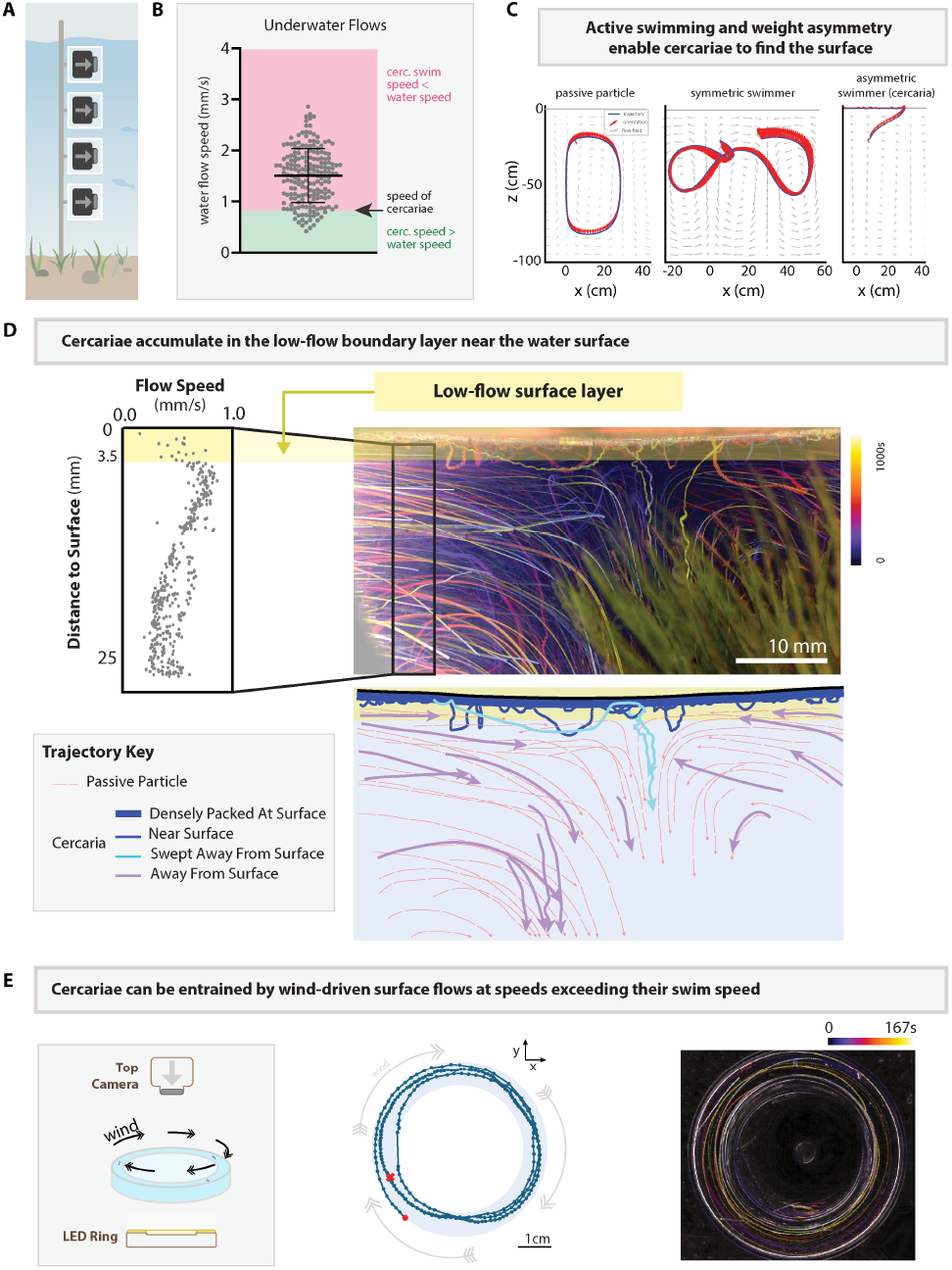
Cercariae rely on their weight asymmetry to navigate flows and find the water surface. **A** Schematic representation of the experimental setup for underwater imaging at the site of Mbakhana, Senegal. **B** Measured speed of the passive particles acting as water flow tracers, relative to the speed of swimming cercariae measured in a laboratory setting. Data measured underwater at the Mbakhana site. **C** Comparison between simulated trajectories of a passive asymmetric particle, a symmetric swimmer and an asymmetric swimmer (cercaria) within a thermal flow cell. **D** Top-left: Flow velocity measurements from a reconstituted natural environment containing water, parasites and vegetation from Mbakhana, revealing a low-flow layer several millimeters thick under the air-water interface. Top-right: overlay of a picture of the aquarium and the temporal color-coded trajectories of passive particles and cercariae. Bottom: schematic representation distinguishing between the trajectories of passive particles and cercariae. **E** Left: schematic representation of the experimental setup to measure the displacement of cercariae exposed to surface wind. Center: Trajectory of a single wind-driven cercaria over 40cm (start: red circle, end: red cross). Right: temporal color coded projection of the trajectory of multiple cercariae exposed to surface wind (view from top).

### Cercariae navigate flows thanks to their weight-asymmetric body structure

Thermal gradients in natural water bodies generate convective circulation patterns that typically entrap passive particles in persistent loops (fig. 5C). Our asymmetric rod model demonstrates that the inherent weight asymmetry between cercarial head and tail functions as a biophysical orientation mechanism, causing the organism to behave as a microscale gravitational pendulum that consistently reorients vertically (fig. 5C). This gravitational reorientation, coupled with active propulsion, guides cercariae toward the surface within the convective field. Quantitative analysis shows this morphological adaptation significantly reduces entrainment in recirculating thermal flows compared to both passive particles and symmetric swimmers (fig. S15). While some cercariae remain captured in high flow areas, the majority find and stay at the water surface (fig. S15).

### Cercariae exploit the surface sublayer to stay near the air-water interface

A crucial discovery of our study is that the air-water interface provides cercariae with a hydrodynamic refuge from bulk water currents. In a reconstituted natural environment with vegetation and cercariae from Mbakhana, Senegal, we observed that cercariae accumulate in a zone of reduced flow velocity (fig. 5D, movie S10) approximately 3.5mm thick at the air-water interface. This distinctive boundary layer, documented in marine ecological studies (*61*), creates a protected zone within the otherwise high flow environment.

Critically, the air-water interface is also characterized by near-zero vertical flow due to the boundary condition of the water surface. This physical constraint combined by the lift force arising from interfacial interaction implies that once cercariae reach the surface, they remain trapped in this two-dimensional plane, preventing them from being swept back into the bulk water volume by vertical currents. It is important to note that this trap is “purely” hydrodynamic and cercaria body do not utilize a hydrophobic patch to attach to air-water interface. This gives the parasite freedom to easily leave the interface by essentially stopping swimming.

By restricting their movements to this sublayer, cercariae transform a three-dimensional volumetric search within complex flows into a more manageable two-dimensional surface exploration within a hydrodynamically protected region. The impact on transmission efficiency is profound: for a surface exploration of 1*m*^2^ with 500 cercariae exploring at 1*mm*^2^/*s*, the required exploration time would be approximately 33 minutes — a three-order-of-magnitude reduction compared to volumetric exploration (23 days). This aligns with both the parasite’s lifespan and typical human water contact duration.

### Cercariae exploit wind-driven surface currents for long-distance dispersal

While surface swimming provides cercariae with a protected microhabitat in the surface layer, where they can remain stable due to the absence of vertical flows, they also leverage wind-driven lateral surface currents for long-distance dispersal. In a ring-shaped aquarium with controlled air-flow (fig. 5E), we tracked cercariae entrained by wind-induced currents at 6 mm/s (fig. S16A, movie S11), which is six times their swimming speed (fig. S16A). This entrainment enabled rapid transport over 40 cm within our experimental setup (fig. 5E, fig. S16B), limited only by apparatus dimensions, suggesting potential for substantially greater distances in natural environments. This passive transport mechanism significantly enhances cercarial dispersal capability, allowing exploitation of meteorological events for extensive spatial distribution without depleting the limited energy reserves of the parasites.

### Discussion and outlook

Our biophysical model reveals that the diverse swimming repertoire stems from cercarial morphology physics rather than neural control. The weight asymmetry between head and tail creates a gravitational pendulum effect that, combined with binary swimming activation, generates all observed behaviors. This exemplifies morphological computation, where physical structure rather than neural complexity solves adaptive challenges. Since cercariae rely on finite glycogen reserves (*62*), offloading computational demands to morphology conserves critical energy, extending host-seeking time. Similar principles appear throughout biology (*63*): numerous organisms leverage physics-based controllers encoded in morphology rather than energy-intensive neural processing for essential functions (*64–66*).

Our physics-based approach shows that cercariae behave as active, weight-asymmetric particles in realistic hydrodynamic flows, allowing us to forecast their distribution. We find that wind-driven currents can entrain cercariae—transporting them at up to six times their swimming speed over distances exceeding 40 cm, which is consistent with theoretical models (*67*) and data on avian schistosomes (*68*). Though marine organisms employ similar gravitactic mechanisms for different purposes (*69*), computational frameworks from marine biology (*70, 71*) could be repurposed to predict zones of increased schistosomiasis risk.

The surface accumulation behavior resolves a central paradox in schistosomiasis transmission: cercariae achieve efficient host-finding despite navigational limitations, low concentrations, and modest propulsion. By leveraging physics dictated by their weight-asymmetric morphology, cercariae exploit fluid dynamics at the air-water interface, transforming an impossible volumetric search into a feasible surface exploration. Our order-of-magnitude calculations show a remarkable reduction in host-encounter time from 23 days to 33 minutes. In reality, both estimates underestimate true search times due to redundant path coverage, but this effect is far more pronounced for volumetric swimmers, which are physically constrained to move laterally only a few millimeters before gravitational reorientation forces vertical movement (fig. S11). Surface swimmers, with their sustained circular trajectories, maintain effective lateral exploration, making the actual efficiency gain likely greater than our three-order-of-magnitude estimate suggests.

In endemic regions, individuals average 180 hours of annual water contact (*72*). Under a volumetric swimming model, a single encounter between a cercaria and a human would require approximately 552 hours (23 days). With only 10% of skin-penetrating cercariae developing into mature schistosomes (*73, 74*), this predicts merely (180/552) × 10% = 0.03 worms per person at risk per year. This is in stark contrast with clinical observations of approximately 100 worms per person at risk (*75*). Surface swimming, however, resolves this discrepancy. With cercariae concentrated at the air-water interface, the calculated time-to-infection decreases dramatically to 33 minutes (0.55 hours), resulting in (180/0.55) × 10% = 33 new worms per person annually, closely aligning with the average measured burden of 100 worms per person given that the worms have a lifetime of several years (*76*). Our work provides a mechanistic explanation for infection rates that volumetric models fundamentally cannot account for, bridging the gap between parasite behavior and epidemiological reality. This highlights how critical surface swimming is for effective infection, but also reveals critical vulnerability in the schistosome life cycle: preventing surface swimming could reduce by 1 − 0.03/33 = 99.9% the number of infection. Interventions targeting the water surface to prevent surface swim, such as biodegradable toxic interfacial films, surfactants such as washing soap (used commonly in the field) that disrupt surface tension, or physical barriers could reduce transmission without the ecological drawbacks of traditional molluscicides (*77*). Combined with mass drug administration, these physics-informed strategies offer a promising, environmentally sustainable approach to schistosomiasis control.

In conclusion, the cercarial body, functioning as a weight-asymmetric “surface-finding machine,” exemplifies how evolution offloads computational demands to physical structure, enabling cercariae to exploit the air-water interface despite ambient flows that would otherwise overwhelm their swimming capacity. Our discovery of surface swimming as the primary infection strategy for schistosome cercariae resolves a longstanding transmission paradox. By shifting from volumetric to surface-constrained exploration, these parasites achieve a thousand-fold improvement in host-finding efficiency without sophisticated neural navigation. At the surface, cercariae benefit from the boundary condition that prevents vertical flows, allowing them to remain stably at this optimal host-encounter zone. Simultaneously, they exploit horizontal wind-driven currents for passive long-distance dispersal at speeds exceeding their swimming capacity. This finding identifies the interface as a previously unrecognized vulnerability in the parasite life cycle, suggesting interventions targeting the water surface could complement traditional control strategies.

## Acknowledgments

We thank all members of the Prakash laboratory (past and present) for helpful discussions. We thank R. Konte for assistance with and valuable feedback on figures. We thank B. Foster for valuable modeling conversations. We thank G. de Leo for useful discussion and sharing his valuable contacts and experience in Senegal. We thank the Station d’Innovation Aquacole for hosting some of the experiments, as well as all the members of the station for their warm welcome. The authors extend their sincere gratitude to the Senegalese communities that graciously welcomed and accommodated our research efforts. The infected snails were provided by the Schistosomiasis Resource Center of the Biomedical Research Institute (Rockville, MD) through NIH-NIAID Contract HHSN272201700014I. NIH: Biomphalaria glabrata (NMRI) exposed to Schistosoma mansoni (NMRI).

## Author contributions

M.P. supervised the project. M.H. and M.P. designed the research project. M.H. developed custom instrumentation, performed the experiments and analyzed the data. A.N. provided training and assisted with snail collection at field sites in Senegal. M.H. developed the asymmetric rod model. I.H. developed dumbbell model with time-varying radius. M.H., I.H. and M.P wrote the manuscript.

## Data and materials availability

Original videos, data and code are available on the Dryad platform (http://datadryad.org/share/gFkdjvAh_kJvFZSFOxizCLpb7JLm7D5HKuaYNKoaGGw).

## Supplementary materials

Materials and Methods

Supplementary Text

Figs. S1 to S15

References (*1-81*)

Movie S1-S11

## Supplementary Materials

### Materials and Methods

#### Harvesting cercariae

*Biomphalaria glabrata* snails infected with *Schistosoma mansoni* were obtained from the Biomedical Research Institute (BRI, Rockville, MD) and maintained according to BRI Standard operating procedures (*78, 79*). Snails were maintained in plastic containers with artificial pond water at 20-22°C with a 24-hour dark cycle. The artificial pond water was prepared by combining four stock solutions: (1) FeCl_3_·6H_2_O (0.25 g/L); (2) CaCl_2_·2H_2_O (12.9 g/L); (3) MgSO_4_·7H_2_O (10 g/L); and (4) a phosphate buffer prepared by dissolving 34 g KH_2_PO_4_ in 500 mL water, adjusting to pH 7.2 with approximately 175 mL 1N NaOH, adding 1.5 g (NH_4_)_2_SO_4_, and bringing the final volume to 1 L. For 20 L of artificial pond water, we combined 10 mL FeCl_3_ solution, 50 mL CaCl_2_ solution, 50 mL MgSO_4_ solution, and 25 mL phosphate buffer, then diluted to 20 L with deionized water. Snails were fed every two days with organic romaine lettuce leaves. Water was changed every 2-3 days. During water changes, snails were carefully transferred using featherweight forceps after being collected on a fine sieve. To obtain cercariae for experiments, individual infected snails were placed in small containers with 3 mL of artificial pond water and exposed to bright light (approximately 1000 lux) for 1 hour to stimulate cercarial shedding. The cercarial suspension was collected afterwards.

#### Cercarial imaging

For high-resolution imaging of cercarial swimming behaviors, two experimental setups were employed. For detailed observations of individual cercariae, specimens were placed in 1×1 cm cross-section polystyrene spectrometry cuvettes containing artificial pond water. For observations under more naturalistic conditions, cercariae were introduced into a 15×15×15 cm glass aquarium containing water, vegetation, and substrate collected from Mbakhana, Senegal, where schistosomiasis is endemic.

Two imaging systems were used: (1) Akaso EK700 cameras equipped with a modified 25 mm tube lens and a 100 mm objective lens, capturing at 30 frames per second; and (2) A Panasonic Lumix GH6 camera with a 40mm/f2.8 TTartisan lens, capturing at up to 300 frames per second. For comprehensive analysis of swimming behaviors, particularly at the air-water interface, dual-camera setups were arranged with two Akaso EK700 cameras to simultaneously capture top and side views (fig. 1C). Illumination was provided by diffuse LED ring lights, each generating 250 lux on the cercariae.

For imaging cercarial responses to potential host cues, a small fragment of naturally shed dead skin (approximately 1×1 mm) donated by one of the authors was introduced in the imaging chamber, and cercarial movements were recorded for 30 minutes before and after introduction of the stimulus.

#### Model

A biophysical model was developed to simulate cercarial swimming behavior based on the physical properties and weight distribution of *S. mansoni* cercariae. The cercaria was represented as a two-part rigid body with asymmetric weight distribution between the head and tail regions. The model incorporated gravitational forces, viscous drag, propulsive forces generated by tail movements, and surface tension effects at the air-water interface. Equations of motion were derived under the assumption of low Reynolds number hydrodynamics (see Supplementary Text). The resulting differential equations were solved numerically using python with the RK45 solver. Simulation parameters were adjusted to match experimental observations of cercarial swimming speeds and trajectories.

#### Data analysis

Cercarial swimming trajectories were extracted from video recordings using the trackmate module (*80*) in ImageJ (*81*), and further analyzed with python. Individual cercariae were tracked over time to quantify swimming speed, orientation angle, and swimming mode transitions. Surface accumulation was quantified by determining the percentage of cercariae within 1 mm of the air-water interface compared to the total number visible in the field of view. For flow field measurements in natural environments, passive tracer particles in the water were tracked using using the trackmate module (*80*) in ImageJ (*81*). Statistical analyses were performed using GraphPad Prism 10.4.1

#### Use of AI-assisted technologies

This manuscript was initially edited with the assistance of Claude 3.7 Sonnet (Anthropic, version 2025), a large language model, to improve language style and clarity, particularly for non-native English speaking authors. The model was used solely as an input for language refinement suggestions and not for data analysis, figure creation, or scientific interpretation. Prompts used included variations of “improve this text; the style should be that of a scientific paper” applied to draft sections. All text generated by Claude was subsequently thoroughly re-edited by the authors. All scientific content, analyses, and conclusions were generated and verified by the authors. All authors reviewed and approved the final text to ensure accuracy and appropriate scientific communication.

### Supplementary Text

#### Cercaria modeled as a weight-asymmetric self-propelled rod

We model the swimmer as a rod with weight asymmetry moving in a viscous fluid with a spatial flow field. The dynamics incorporate hydrodynamic interactions, gravitational effects, and boundary conditions at the surface.

- *L*: Length of the rod.
- *W*: Width of the rod.
- *µ*: Fluid viscosity.
- (*x, z, θ*): Position and orientation of the rod, where *θ* is the angle relative to the horizontal.
- *F*_*p*_: Self-propelling force along the rod’s axis.
- *F*_*g*_: Gravitational force.
- *T*_*g*0_: Gravitational torque due to weight asymmetry.
- *k*_*s*_: Contact stiffness with the boundary.

At low Reynolds number, the relationship between forces/torques and velocities/angular velocities is linear and described by a mobility matrix **M**. For a slender rod with length *L* and width *W* in a fluid with viscosity *µ*, the mobility matrix is given by:

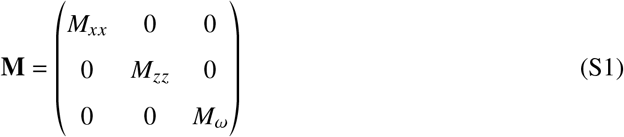

Where:

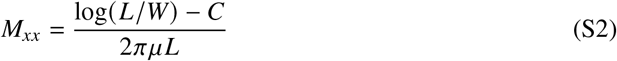

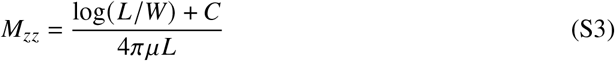

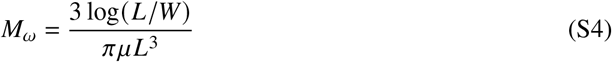

Here, *C* = 0.5 is an empirical correction factor. This matrix relates the forces and torques to the resulting velocities in the rod’s reference frame.

The model can accommodate any arbitrary ambient flow field. While the framework is general, we use thermal convection cells (Rayleigh-Bénard) as an illustrative example. The flow field is modeled using a stream function approach, producing a spatially varying flow with boundary layer effects:

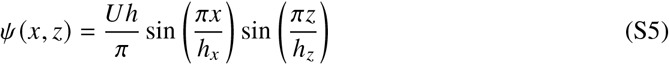

From this stream function, the velocity components are derived:

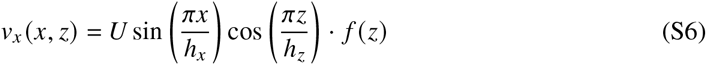

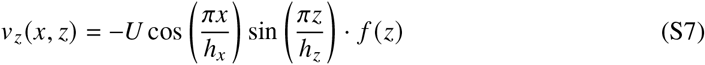

Where *U* is the characteristic flow velocity amplitude, and *f* (*z*) = min(|*z*|/*δ*, 1) is a boundary layer factor that scales the flow velocity near the boundary, with *δ* being the boundary layer thickness. The local vorticity in the flow field is:

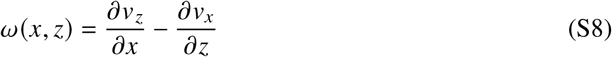

It should be noted that this specific flow pattern represents one possible scenario, and the model formulation allows for the implementation of various other flow fields to study different environmental conditions.

The orientation dynamics follow a Jeffery’s orbit, taking into account the local flow vorticity and the aspect ratio of the rod:

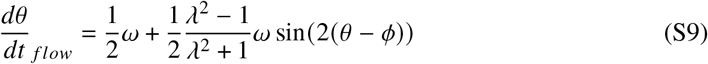

Where *λ* = *L*/*W* is the aspect ratio, *ω* is the local vorticity, and *Φ* = arctan 2(*v*_*z*_, *v*_*x*_) is the direction of the local flow.

The forces acting on the rod include:

1. Self-propelling force *F*_*p*_

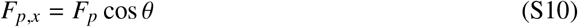

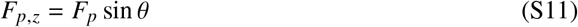
2. Gravitational force *F*_*g*_ acting downward in the *z* direction.
3. Contact force with the boundary when *z* > 0, acting in the z direction:

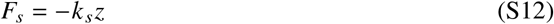
4. The associated contact torque:

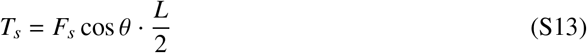
5. Gravitational torque due to weight asymmetry:

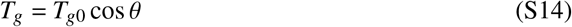

The system’s equations of motion combine the contributions from direct forces and torques through the mobility matrix and the influence of the ambient flow:

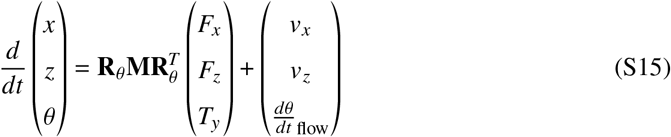

Where **R**_*θ*_ is the rotation matrix that transforms from the rod’s frame to the lab frame:

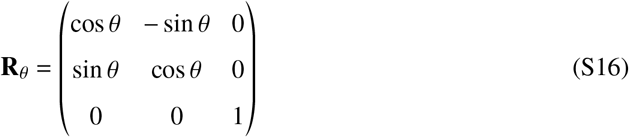

This system of coupled ordinary differential equations can be solved numerically using standard methods such as adaptive time-stepping methods like those implemented in the *solve iv* function from the *SciPy* python library.

#### Cercaria modeled as two oscillating spheres connected by a dragless rod

The interaction of microswimmers with surfaces has been extensively studied, revealing how organisms at low Reynolds number (*Re*) interact hydrodynamically with nearby boundaries. Analytical and numerical studies have characterized how swimmers are attracted, repelled, or reoriented by planar surfaces. Models ranging from simplified squirmers and force multipoles (*43–47*) to detailed shapes like sheets, rods, and linkage-based swimmers (*48–52*) have been used to represent biological swimmers. Near a stress-free interface, the complexity increases significantly (*38, 53–58*), especially when interface deformation—characterized by the capillary number (*Ca*)—becomes important.

Schistosome cercariae swim at low Reynolds numbers 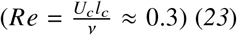, where viscous forces dominate inertial forces. We define the characteristic length *l*_*c*_ = *L*/2, where *L* = 500, *µ*m is the cercarial body length. The characteristic time is *t*_*c*_ = 1/ *f* where *f* = 10Hz is the typical beat frequency. *U*_*c*_ is its characteristic swimming speed. Cercariae utilize elastohydrodynamic coupling in their ‘T-swimmer’ gait to break time-reversal symmetry (*23*). In our observations, cercariae swim close to the air-water interface (*h*/*l*_*c*_ ≈ 0.2), with negligible deformation 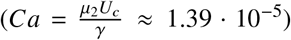. Under these conditions, lubrication effects are crucial (*59*), and far-field approximations (*38, 53, 54, 56*) fail to accurately capture the dynamics. Given the small capillary number, interface deformation can be neglected (*26, 27, 59*), allowing a purely planar interface assumption.

##### Kinematic Model

We model cercariae as two spheres connected by a massless, dragless rod in 2D Stokes flow (*23, 37*). The velocities of each sphere 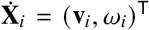 where *i* = 1, 2 for each sphere are related to the swimmer’s centroid velocity **V**_*b*_ = (**v**_*b*_, *ω*_*b*_)^T^ via

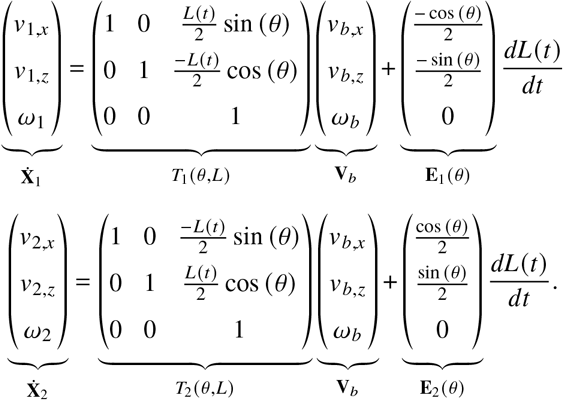

Concatenating both spheres gives,

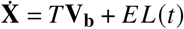

where **X** = (**X**_**1**_, **X**_**2**_)^T^, *T* = (*T*_1_, *T*_2_)^T^ and *E* = (*E*_1_, *E*_2_)^T^. Since the linearity of Stokes flow implies,

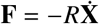

where *R* is the grand-resistance matrix, the force on each sphere in terms of body motion is then,

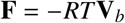

where **F**_*i*_ = (**f**_*i*_, *m*_*i*_)^*T*^. Likewise it can also be shown that kinematic equations imply,

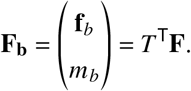

As a result, the total force on the body is,

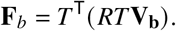

For some external body force on the two sphere system, **F**_ex_ = (0, *F*_*g*_, *T*_*g*_ cos (*θ*)), the force free condition in Stokes flow gives,

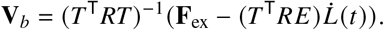

We nondimensionalize the differential equations with length scale *l*_*c*_ and time scale *t*_*c*_ and integrate with Euler stepping Δ*t* = 0.001 in MATLAB.

##### Resistance of a sphere near a planar interface

The Grand resistance matrix *R* is block-diagonal,

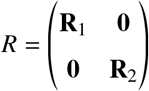

where the friction coefficients are both a function of each sphere’s radius *a* and separation to the interface *h*,

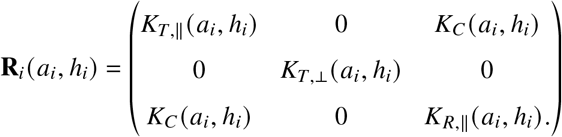

Fig. S12 shows the dimensionless resistance values for a single sphere as a function of the separation from the interface. These friction coefficients are fit to numerical solutions (*27,29*) using global interpolation functions (*28*),

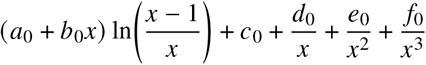

where *x* = *h*/*a*.

In the present formulation we retain only the *single–particle* wall corrections to the resistance of each sphere. Consequently we neglect (i) Faxén–type finite–size corrections (*30*); (ii) hydrodynamic interactions between the two real spheres, their first images and the infinite hierarchy of reflections, which has been well documented for wall–Stokesian-dynamics pair mobilities (*31–33*). To obtain more accurate trajectories and interface-induced reorientation, these omitted contributions should be incorporated.

##### Soft repulsion from the interface

To prevent the dumbbell swimmer from penetrating the interface using time steps of size Δ*t* = 0.001, a soft steric exclusion from the interface (*z* = 0) is enforced through a steep power-law. This approach is biologically motivated: Schistosoma cercariae are known to be hydrophilic and remain submerged in the aqueous phase, avoiding air–water contact. The soft repulsion thus captures the physical constraint that their bodies do not breach the surface tension barrier at the interface. Letting *δ*_*i*_ = max(10^−10^, −*z*_*i*_ − *a*_*i*_) denote the instantaneous wall–surface gap of sphere *i* = 1, 2, the wall-normal repulsive force is prescribed as an exponentially decaying repulsion

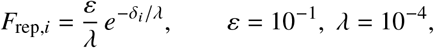

and is applied along 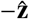. The two forces are assembled in the laboratory frame as

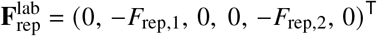

and rotated into the swimmer frame via 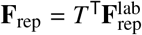 before being added to the rigid-body force balance.

##### Stroboscopic map and phase portrait

For every beat cycle of duration *T* = 1/ *f* we integrate the full hydrodynamic model and record the swimmer’s vertical position *z* and pitch angle *θ* at the cycle’s end (since the interface is infinite, the equations are invariant under shifts in *x*). This yields the Poincaré map

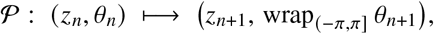

whose trajectories build the (*z, θ*) phase portrait in fig. 3D–E of the main text.

##### Linear stability of the fixed point

A fixed point satisfies *P*(**s**^∗^) = **s**^∗^ with

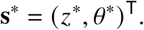

Local dynamics are governed by the Jacobian

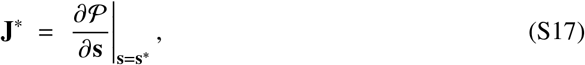

assembled numerically with centred differences. Its eigenvalues *λ*_1,2_ and eigenvectors **v**_1,2_ describe the linear response of the stroboscopic Poincaré map: the fixed point is linearly stable whenever |*λ*_1,2_| < 1. For the two parameter sets investigated (*T*_*g*_ = 0 and *T*_*g*_ = 0.15) the eigenvalues are numerically

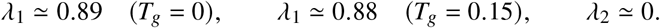

These *λ*_*i*_ are Floquet multipliers of the one-period map *P*, hence they quantify how an infinitesimal perturbation evolves over a single actuation cycle: a component along **v**_*i*_ is simply rescaled by *λ*_*i*_. Consequently, |*λ*_1_| ≈ 0.88–0.89 corresponds to some fractional contraction per cycle in the slowest stable direction, whereas *λ*_2_ ≃ 0 shows that perturbations along **v**_2_ are effectively annihilated within one cycle, confirming that the fixed point is hyperbolically attracting.

##### Fixed-point search

Fixed points are the roots of the nonlinear system

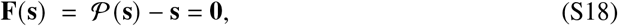

where **s** = (*z, θ*)^T^. We first lay down a coarse grid in the (*z, θ*)-plane where ach grid node serves as an initial guess for a Newton–Raphson iteration on (S18). Converged roots are clustered with a tolerance 10^−3^ to remove duplicates, yielding attracting fixed points and one saddle (for *T*_*g*_ = 0).

##### Separatrix identification

(*T*_*g*_ = 0) A coarse grid search in the (*z, θ*) plane locates a unique saddle, diagnosed by one Floquet multiplier |*λ*_st_| ≃ 0.73 < 1 and one |*λ*_un_| ≃ 1.60 > 1. Let the corresponding eigenvectors be **v**_st_ and **v**_un_. From the saddle position **s**_sad_ we seed four nearby points

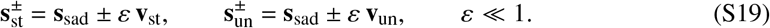

Iterating 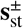 backwards with the inverse Poincaré map traces the two stable branches converging onto the saddle. Iterating 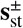 forwards with the Poincaré map traces the two unstable branches departing from the saddle. Together, the branches form the separatrix that divides the phase space into regions of distinct long-time behavior.

##### Basin stability

Following the probabilistic framework introduced in (*25*), the Poincaré map is iterated to assess the global robustness of the surface-swimming fixed point **s**^∗^ = (*y*^∗^, *θ*^∗^)^T^. The Monte-Carlo domain of admissible initial conditions is

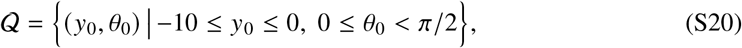

further restricted by a no-penetration screen that rejects any (*y*_0_, *θ*_0_) for which either sphere would start above the interface. Basin stability is estimated from *N* = 10^3^ uniformly drawn samples,

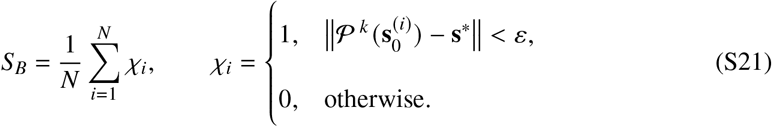

with *ε* = 10^−3^ and at most *k*_max_ = 600 strobes per trajectory. The statistical uncertainty is

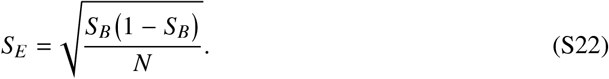

Fig. S13 plots the resulting estimates of *S*_*B*_ (with error bars *S*_*E*_) as a function of the nondimensional buoyant torque *T*_*g*_, revealing how increased torque progressively shrinks the basin of attraction.

##### Simulation parameters

The simulation parameters for main text fig. 3 are nondimensionalized by characteristic length (*l*_*c*_), time (*t*_*c*_), force (*F*_*c*_) and torque (*τ*_*c*_) scales,

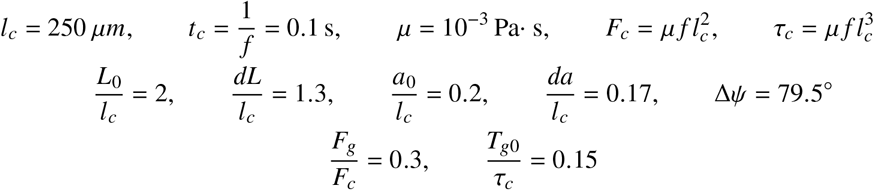

For the dumbbell model, the gravitational force *F*_*g*_ on the cercariae is calculated using an estimated cercarial density and volume: *F*_*g*_ = (*ρ*_*f*_ − *ρ*_*c*_)*V*_*c*_*g*, where *g* is gravitational acceleration, *ρ*_*f*_ and *ρ*_*c*_ are densities of water and cercariae respectively, and *V*_*c*_ is the cercarial volume (*23*). The buoyant torque *T*_*g*0_ is assumed constant, acting at the dumbbell’s centroid, and its magnitude is given by 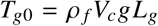, with *L*_*g*_ being the experimentally determined separation between the center of buoyancy (COB) and the center of gravity (COG) (*23*). To constrain the model, we fit a set of dimensionless parameters: *L*_0_, *δL, a*_0_, *δa*, Δ*ψ*. The baseline radius *a*_0_ is selected so that passive sedimentation of a vertically oriented dumbbell (*a*_1_ = *a*_2_ = *a*_0_, *δL* = *δa* = 0) matches the experimentally observed sinking speed of 0.1, mm,s^−1^ (*23*). The length *L*_0_ is chosen to be the body length of the cercariae. Finally, parameters *δL, δa*, and Δ*ψ* are fitted to reproduce the instantaneous tangential swimming velocity observed during horizontal swimming. The validity of these parameters is confirmed by comparing upward swimming velocities from simulations against the experimentally measured tail-first swimming speed of 0.7, mm,s^−1^. Typically, the relative phase offset *δψ* between the two oscillating sphere radii *a*_1_(*t*) and *a*_2_(*t*) can take arbitrary values. The resulting period-averaged dipole strength *p* ∼ cos (*δψ*) sin (Δ*ψ*) determines whether the swimmer behaves as a pusher (*p* > 0) or a puller (*p* < 0) (*34*). Motivated by experimental observations (particle image velocimetry) indicating that cercariae exhibit both pusher-like and puller-like behavior during distinct phases of the beat cycle (power and recovery stroke) (*23*), we approximate *δψ* = *π*/2, yielding a net dipole strength of *p* = 0 over one actuation period.

**Figure S1:**
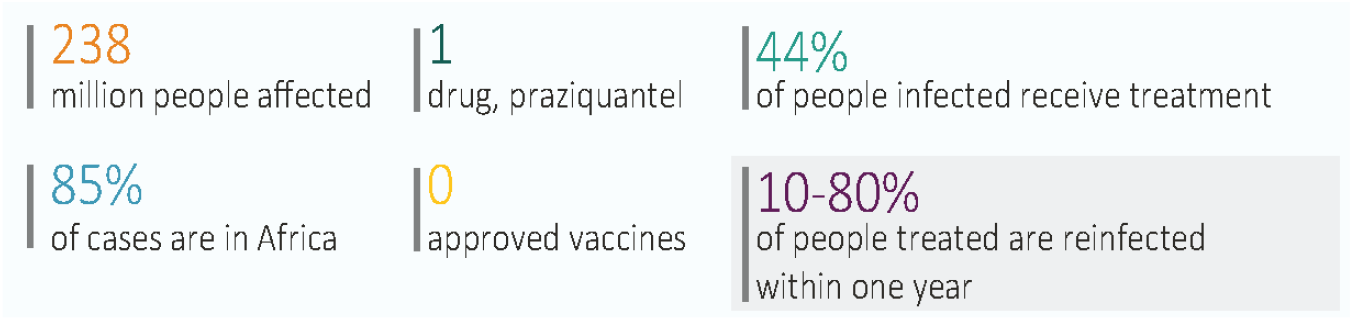
Key numbers on schistosomiasis.

**Figure S2:**
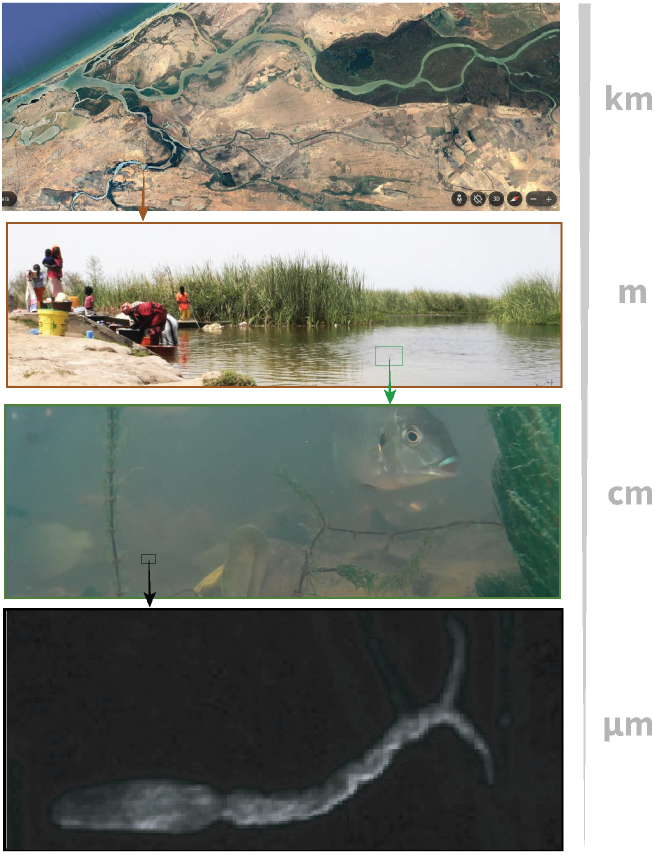
Schistosomiasis is a multi-scale problem. From top to bottom: Satellite picture of the Senegal river (from *Google earth*); picture of the site of Ndiawdoune, Senegal; underwater imaging in the site of Mbakhana, Senegal; picture of a cercaria

**Figure S3:**
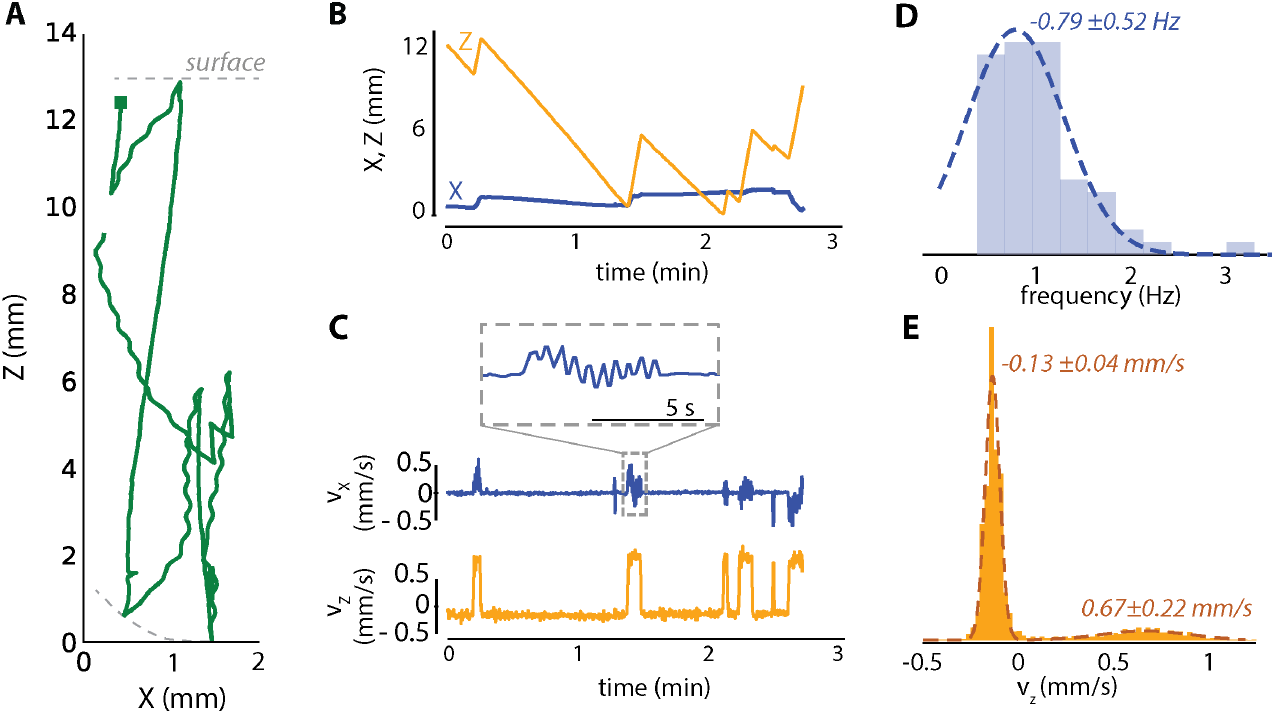
2D trajectory of a cercaria in a thin vertical cuvette. **A** Displacement of a single representative *S. Mansoni* cercaria. A green square represents the starting position. the dotted line represents the bottom of the cuvette and the water surface **B** Evolution over time of the lateral (X) and vertical (Z) position of the cercaria presented in A. **C** Lateral *v*_*x*_ and vertical *v*_*z*_ velocity of the cercaria presented in A. **D, E** Statistical analysis on 17 cercariae. **D** Distribution (blue bars) of the temporal frequency of the spiral motion during the active upward swimming mode, with a fit (dotted line) for a gaussian distribution. The average and standard deviation are noted next to the peak. **E** Distribution (yellow bars)of the instantaneous verrtical velocity over time for cercariae, with a fit (dotted line) for a gaussian bimodal distribution. The average and the standard deviation of each peak (representing the sinking mode and the active swimming upward mode) are indicated next to the peak.

**Figure S4:**
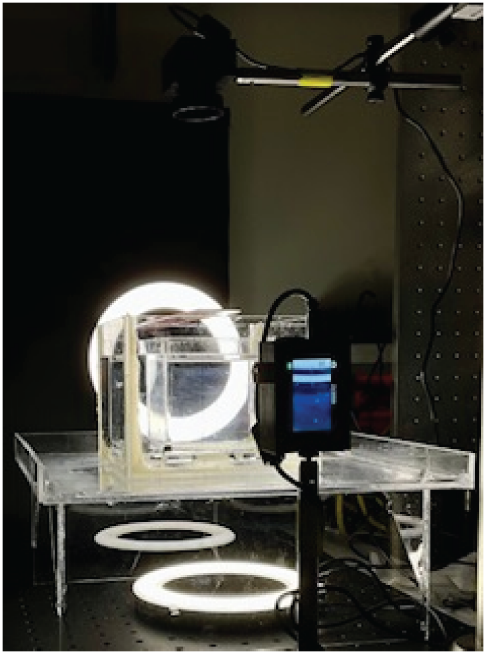
Picture of the experimental setup for 3D imaging. One camera is mounted vertically, the other horizontally. A third camera can be mounted horizontally along the third axis

**Figure S5:**
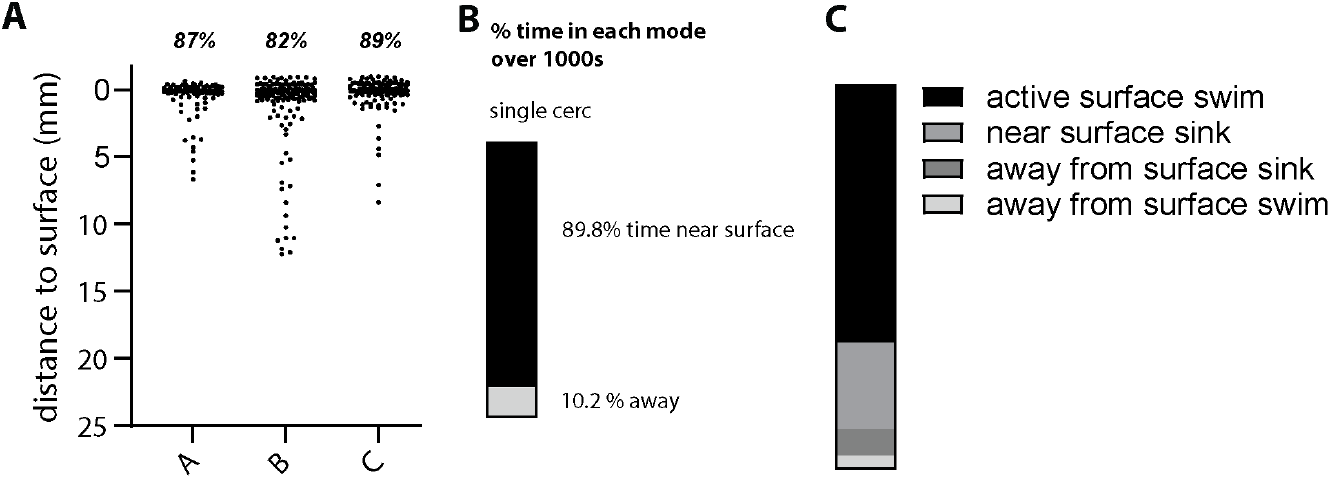
Cercariae accumulate near the air-water interface. **A** Distribution of cercariae in a 1×1×4cm cuvette, in triplicate (biological replicate). The percentage of cercariae within the top 1mm of the water is indicated for each. **B** Distrubution of the time spent near the surface (within 1mm) and away from the surface when tracking a single cercaria over 1000 seconds. **C** Distribution of swimming modes for 100 cercariae.

**Figure S6:**
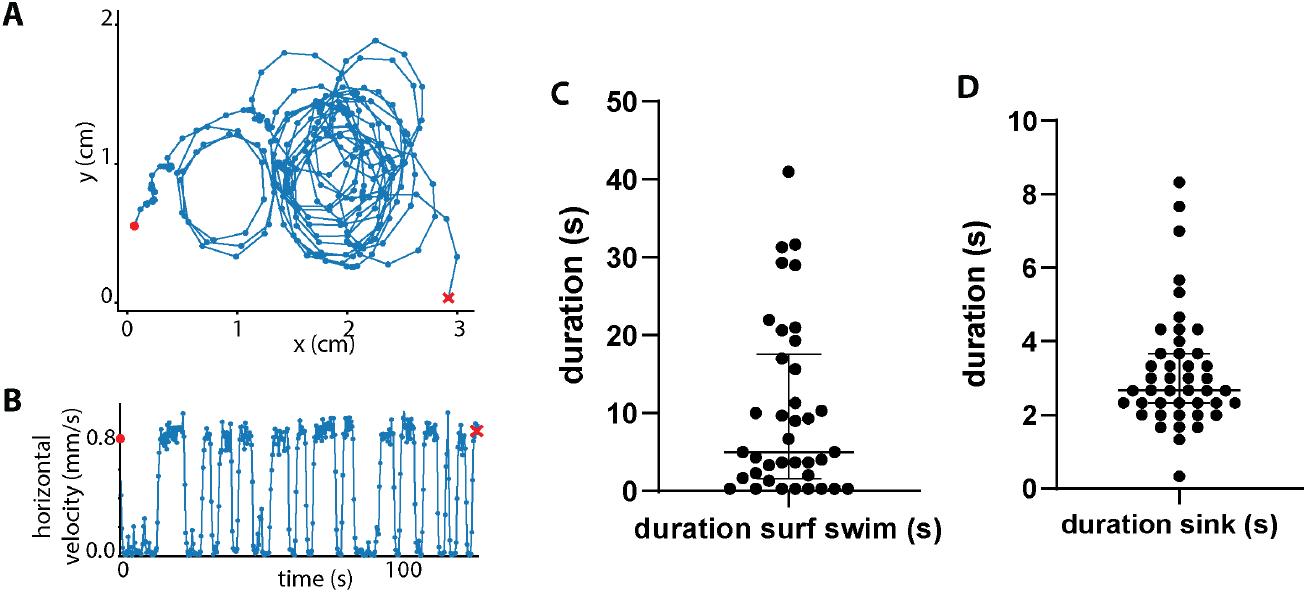
Surface-swimming cercariae alternate swimming with brief periods of sinking. **A** Trajectory of a single cercaria at the water surface, starting at the red dot and finishing at the red cross. **B** Corresponding graph of the horizontal velocity over time. The times with near zero horizontal velocity correspond to sinking events. **C** Distribution of the duration of the surface swim phases. **D** Distribution of the duration of the sinking phases.

**Figure S7:**
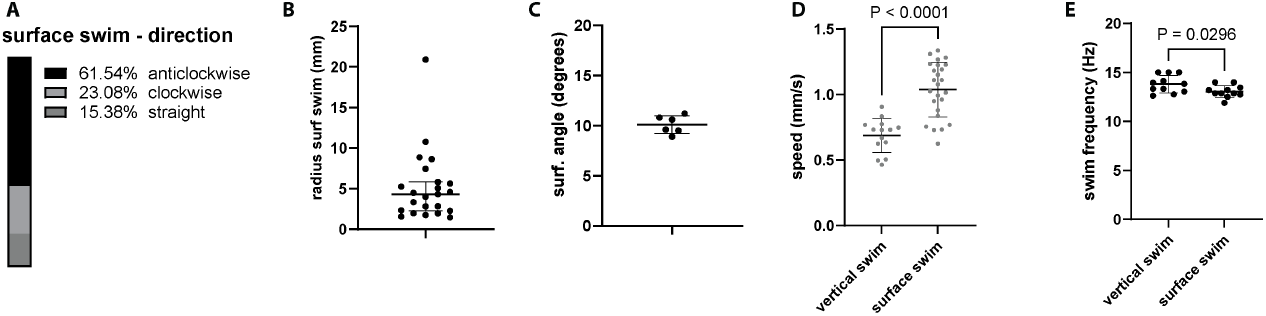
Properties of surface swim. **A** Distribution of the direction of the surface swim (clockwise, anticlockwise or straight) **B** Distribution of the curvature of the circle-like trajectories of surface-swimming cercariae. **C** Distribution of the angle between a surface-swimming cercaria and the water surface. **D** Comparison between the distribution of the swimming speed during vertical swim upwards and during surface swimming. Welch’s two-tailed t-test (unpaired samples, unequal variance, approximately normal distribution, testing for difference in either direction). **E** Comparison between the swim frequency during vertical swim upwards and during surface swim. Welch’s two-tailed t-test.

**Figure S8:**
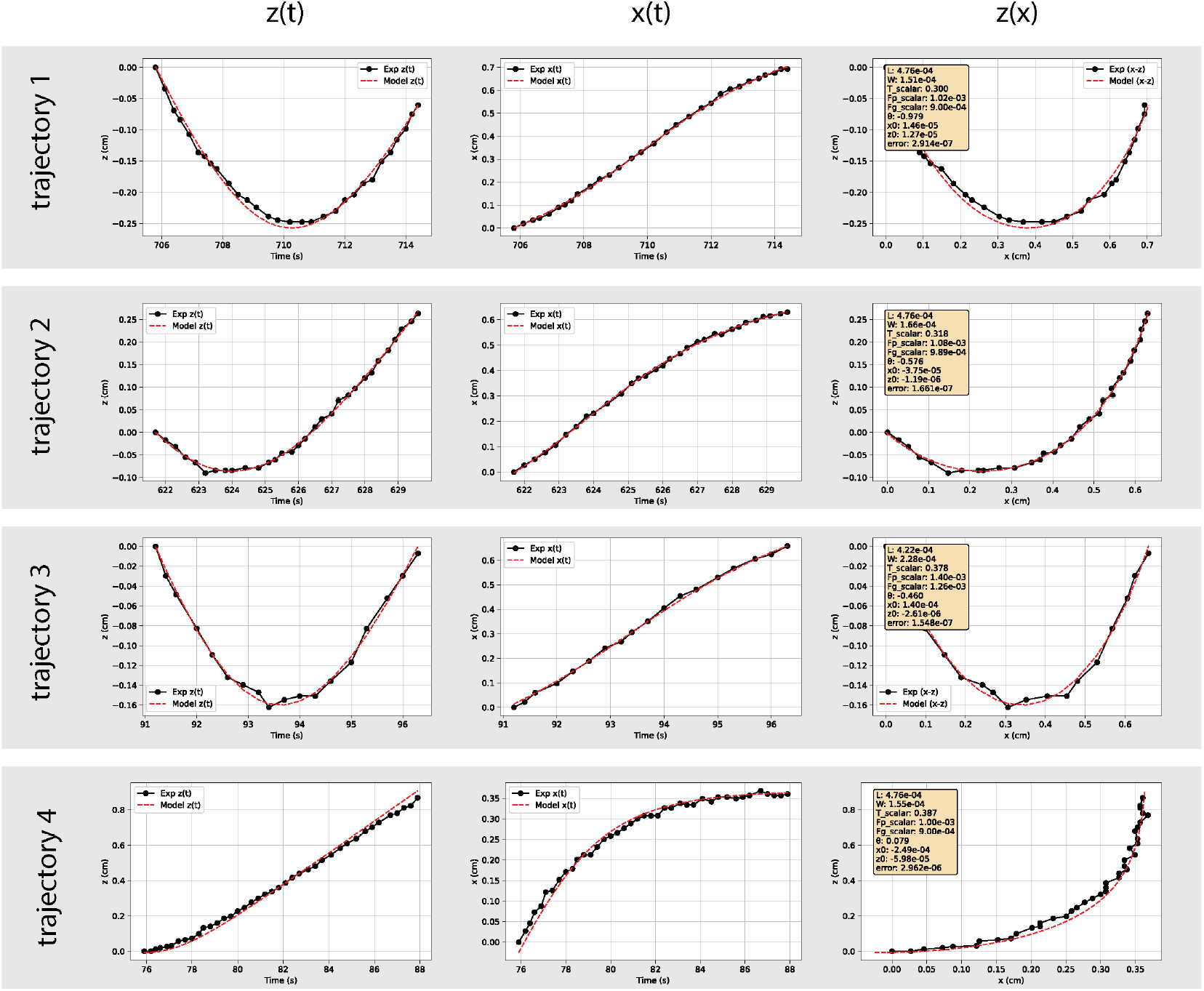
Fit of representative experimental datasets of u-swim with the asymmetric rod model. The experimental data points are represented are black dots, the model as a red line. The fit parameters are indicated in the yellow rectangle for each dataset.

**Figure S9:**
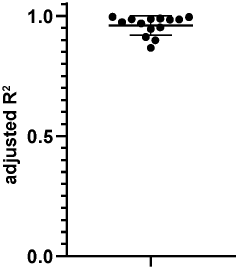
Evaluation of the quality of the fit of the asymmetric rod model. Distribution of the adjusted *R*^2^ parameter for 6 fit parameters and N=14 experimental u-swim datasets.

**Figure S10:**
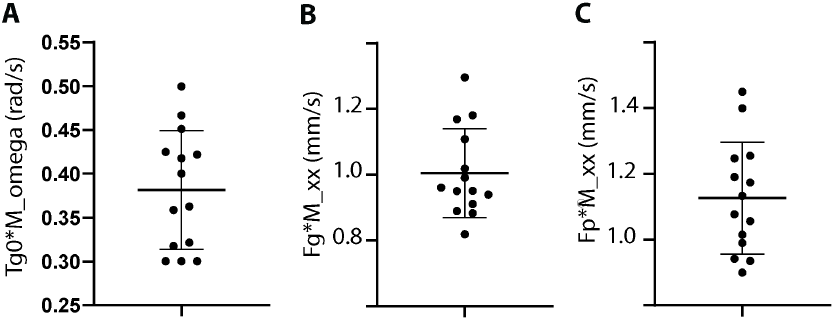
Fitted experimental parameters for the asymmetric rod model. Distribution of the three parameters for the asymmetric rod model: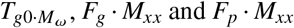, obtained from fitting N=14 experimental u-swim datasets.

**Figure S11:**
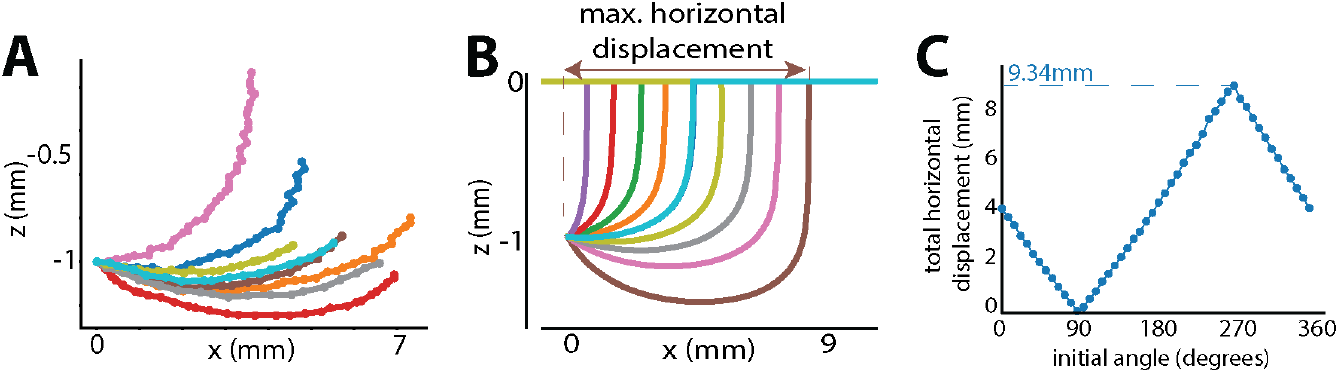
The u-swim trajectory depends only on the initial angle, and the lateral displacement is limited to less than a centimeter. **A** Measured u-swim trajectories as a function of the initial angle. The initial position has been normalized to (−1,0) to facilitate comparison. Each curve has been associated with a random color for clarity. **B** simulated u-swim trajectories as a function of the initial angle. The maximum horizontal displacement achievable is 9.3mm, obtained for an angle relative to the surface of −*π*/2. **C** Calculated horizontal displacement as a function of the starting angle.

**Figure S12:**
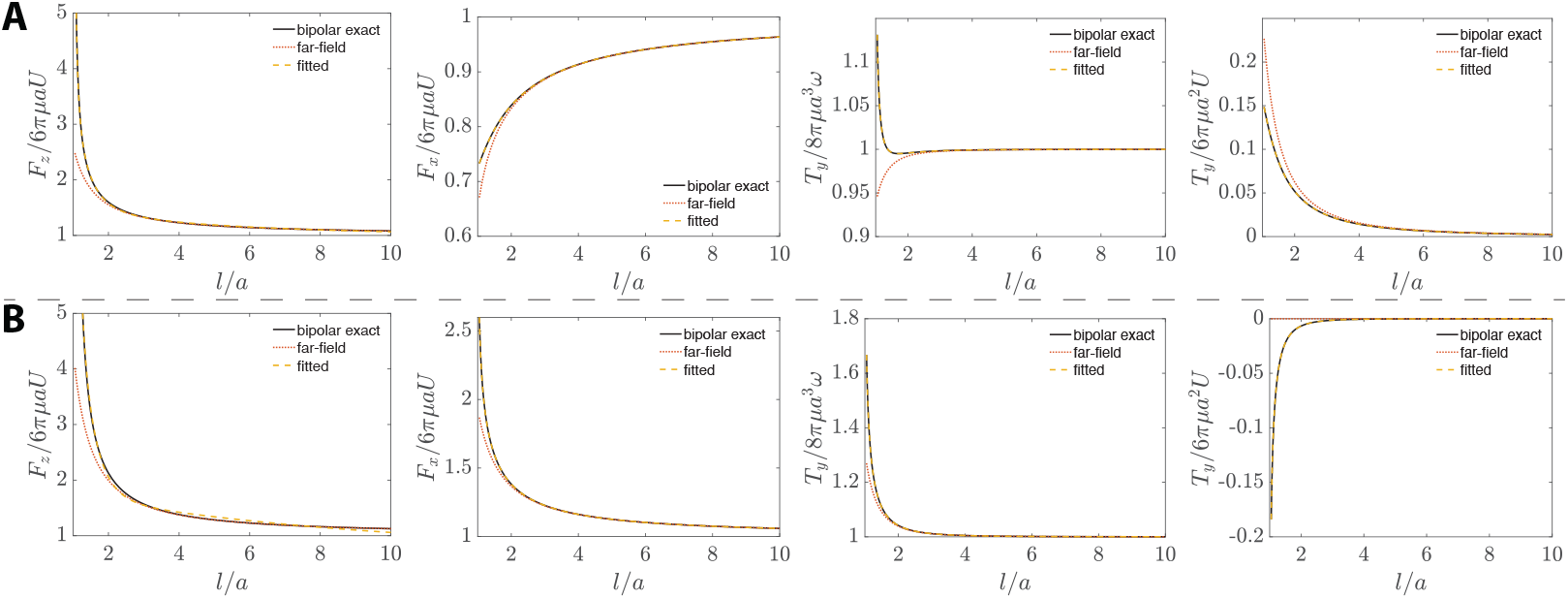
Dimensionless drag and torque on a sphere versus scaled distance to interface. **A** *µ*_1_/*µ*_2_ = 0 in order of *K*_*T*,⊥_, *K*_*T*,∥_, *K*_*R*,∥_ and *K*_*C*_. The far-field solution in (*26*) is compared to the exact bipolar solution (*27, 29*). The exact solution is fitted to a global interpolation formula that is valid in both limits (lubrication and far-field) (*28*). **B** *µ*_1_/*µ*_2_ = ∞ in order of *K*_*T*,⊥_, *K*_*T*,∥_, *K*_*R*,∥_ and *K*_*C*_.

**Figure S13:**
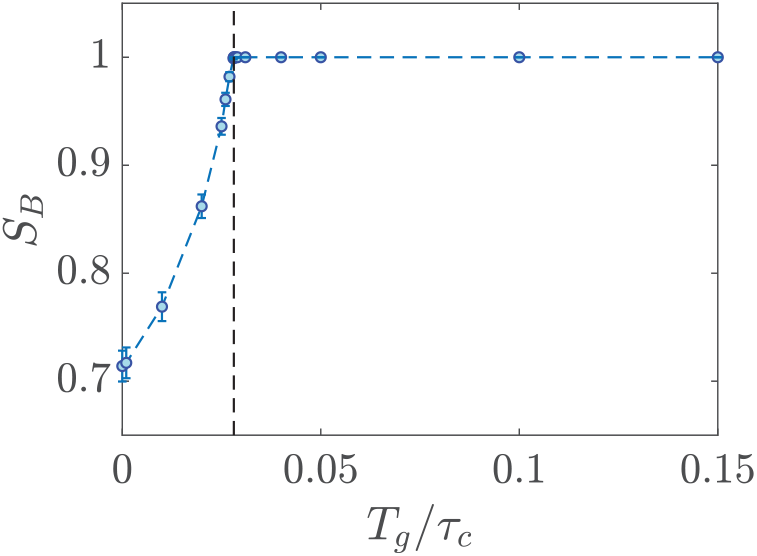
Basin stability versus dimensionless buoyant torque. Error-bars represents ± one standard deviation. *S*_*B*_ = 1 corresponds to the elimination the separatrix, first occurring at *T*_*g*_/*τ*_*c*_ ≈ 0.281.

**Figure S14:**
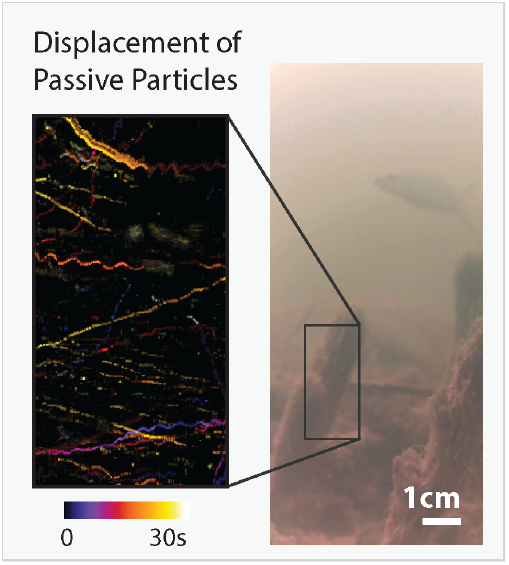
Determination of the underwater flows in a field site using the trajectories of passive particles. see Movie S9

**Figure S15:**
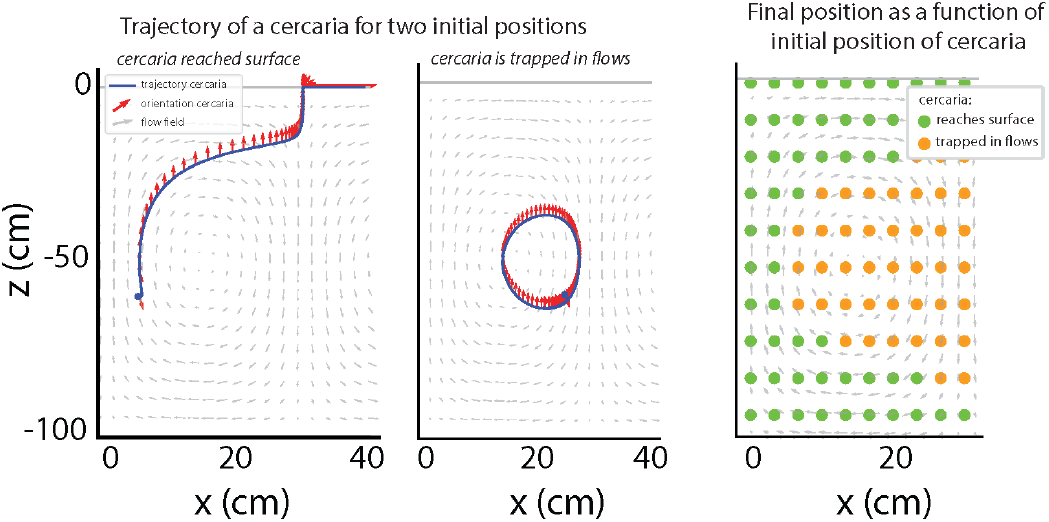
Ability to reach the surface as a function of the initial position within a thermal flow cell.

**Figure S16:**
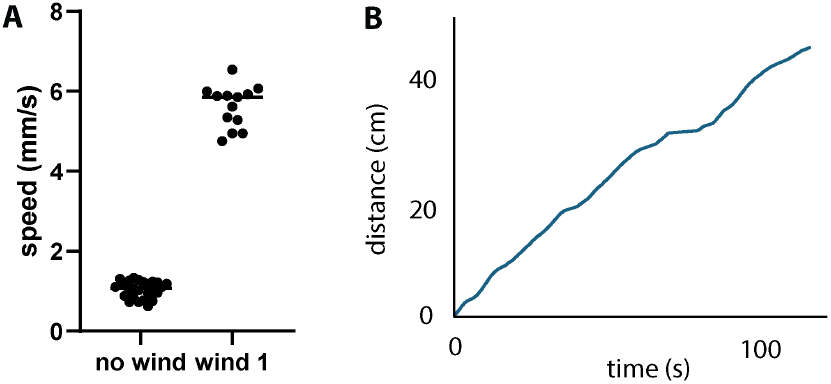
Wind-driven displacement of a cercaria. **A** Cercariae are able to be entrained by the wind at a speed exceeding by several folds their swimming speed. **B** Entrainment by the wind of a cercaria over more than 40cm.

**Caption for Movie S1. Schistosomisis transmission is a multiscale problem**

**Caption for Movie S2. 3D imaging of swimming cercariae**. Animated version of fig. 1C.

**Caption for Movie S4. Cercariae accumulate near the water surface in a realistic environment**. Animated version of fig. 4A.

**Caption for Movie S5. Cercariae accumulate near the water surface in a cuvette**. Horizontal imaging from the side with cercariae in a spectrometry cuvette.

**Caption for Movie S6. Surface-swimming cercariae move in circles - seen from the side** Imaging of a single cercaria swimming at the air water interface. The camera is inclined relative to the horizontal.

**Caption for Movie S7. Surface-swimming cercariae move in circles - seen from the top** Imaging of a cercariae swimming at the air water interface. The camera is imaging the water surface, from the top. Animated version of Figure 4B.

**Caption for Movie S8. High resolution movie of a surface swimming cercaria, seen from the side**. Animated version of fig. 4D.

**Caption for Movie S9. High resolution movie of a surface swimming cercaria, seen from above**. Animated version of fig. 4C.

**Caption for Movie S10. Cercariae are sheltered from water flows when they stay near the water surface** Animated version of fig. 5C.

**Caption for Movie S11. Passive particles act as tracers to measure underwater flows in Mbakhana, Senegal** Animated version of fig. S14.

**Caption for Movie S12. Surface swimming cercariae are entrained by wind-driven surface flows** Animated version of fig. 5D.

## References and Notes

1. Theo Vos, Abraham D Flaxman, Mohsen Naghavi, Rafael Lozano, Catherine Michaud, Majid Ezzati, Kenji Shibuya, Joshua A Salomon, Safa Abdalla, Victor Aboyans, et al. Years lived with disability (ylds) for 1160 sequelae of 289 diseases and injuries 1990–2010: a systematic analysis for the global burden of disease study 2010. The lancet, 380(9859):2163–2196, 2012.

2. Charles H King, Katherine Dickman, and Daniel J Tisch. Reassessment of the cost of chronic helmintic infection: a meta-analysis of disability-related outcomes in endemic schistosomiasis. The Lancet, 365(9470):1561–1569, April 2005.

3. Birgitte Bruun, Jens Aagaard-Hansen, and Susan Watts. The social context of schistosomiasis and its control An introduction and annotated bibliography. Strategies, page 227, 2008. ISBN: 978 92 4 159718 0.

4. Anthony Danso-Appiah, Piero L. Olliaro, Sarah Donegan, David Sinclair, and Jürg Utzinger. Drugs for treating Schistosoma mansoni infection. Cochrane Database of Systematic Reviews, 2013(2), February 2013. Publisher: John Wiley and Sons Ltd.

5. D Engels, L Chitsulo, A Montresor, and L Savioli. The global epidemiological situation of schistosomiasis and new approaches to control and research. Acta Tropica, 82(2):139–146, May 2002.

6. Daniel G. Colley, Amaya L. Bustinduy, W. Evan Secor, and Charles H. King. Human schis-tosomiasis. In The Lancet, volume 383, pages 2253–2264. Lancet Publishing Group, 2014. Issue: 9936 ISSN: 1474547X.

7. J. Meyer-Lassen, A.A. Daffalla, and H. Madsen. Evaluation of focal mollusciciding in the Rahad Irrigation Scheme, Sudan. Acta Tropica, 58(3-4):229–241, December 1994.

8. Christopher M. Hoover, Susanne H. Sokolow, Jonas Kemp, James N. Sanchirico, Andrea J. Lund, Isabel J. Jones, Tyler Higginson, Gilles Riveau, Amit Savaya, Shawn Coyle, Chelsea L. Wood, Fiorenza Micheli, Renato Casagrandi, Lorenzo Mari, Marino Gatto, Andrea Rinaldo, Javier Perez-Saez, Jason R. Rohr, Amir Sagi, Justin V. Remais, and Giulio A. De Leo. Modelled effects of prawn aquaculture on poverty alleviation and schistosomiasis control. Nature Sustainability, 2(7):611–620, July 2019.

9. Susanne H Sokolow, Isabel J Jones, Merlijn Jocque, Diana La, Olivia Cords, Anika Knight, Andrea Lund, Chelsea L Wood, Kevin D Lafferty, Christopher M Hoover, Phillip A Collender, Justin Remais, David Lopez-Carr, Jonathan Fisk, Armand M Kuris, and Giulio A De Leo. Water, dams, and prawns: novel ecological solutions for the control and elimination of schistosomiasis. The Lancet, 389:S20, April 2017.

10. Jason R. Rohr, Alexandra Sack, Sidy Bakhoum, Christopher B. Barrett, David Lopez-Carr, Andrew J. Chamberlin, David J. Civitello, Cledor Diatta, Molly J. Doruska, Giulio A. De Leo, Christopher J. E. Haggerty, Isabel J. Jones, Nicolas Jouanard, Andrea J. Lund, Amadou T. Ly, Raphael A. Ndione, Justin V. Remais, Gilles Riveau, Anne-Marie Schacht, Momy Seck, Simon Senghor, Susanne H. Sokolow, and Caitlin Wolfe. A planetary health innovation for disease, food and water challenges in Africa. Nature, 619(7971):782–787, July 2023.

11. Tayo Alex Adekiya, Raphael Taiwo Aruleba, Babatunji Emmanuel Oyinloye, Kazeem Oare Okosun, and Abidemi Paul Kappo. The Effect of Climate Change and the Snail-Schistosome Cycle in Transmission and Bio-Control of Schistosomiasis in Sub-Saharan Africa. International Journal of Environmental Research and Public Health, 17(1):181, December 2019.

12. Susanne H. Sokolow, Isabel J. Jones, Merlijn Jocque, Diana La, Olivia Cords, Anika Knight, Andrea Lund, Chelsea L. Wood, Kevin D. Lafferty, Christopher M. Hoover, Phillip A. Collender, Justin V. Remais, David Lopez-Carr, Jonathan Fisk, Armand M. Kuris, and Giulio A. De Leo. Nearly 400 million people are at higher risk of schistosomiasis because dams block the migration of snail-eating river prawns. Philosophical Transactions of the Royal Society B: Biological Sciences, 372(1722):20160127, June 2017.

13. Peter Steinmann, Jennifer Keiser, Robert Bos, Marcel Tanner, and Jürg Utzinger. Schistoso-miasis and water resources development: systematic review, meta-analysis, and estimates of people at risk. The Lancet Infectious Diseases, 6(7):411–425, July 2006.

14. Jérôme Boissier, Sébastien Grech-Angelini, Bonnie L Webster, Jean-François Allienne, Tine Huyse, Santiago Mas-Coma, Eve Toulza, Hélüne Barré-Cardi, David Rollinson, Julien Kincaid-Smith, Ana Oleaga, Richard Galinier, Joséphine Foata, Anne Rognon, Antoine Berry, Gabriel Mouahid, Rémy Henneron, Hélene Moné, Harold Noel, and Guillaume Mitta. Outbreak of urogenital schistosomiasis in Corsica (France): an epidemiological case study. The Lancet Infectious Diseases, 16(8):971–979, August 2016.

15. Bruno Gryseels, Katja Polman, Jan Clerinx, and Luc Kestens. Human schistosomiasis. The Lancet, 368(9541):1106–1118, September 2006.

16. Matthew S. Tucker, Laksiri B. Karunaratne, Fred A. Lewis, Tori C. Freitas, Yung san Liang, Allen G.P. Ross, Paul B. Bartley, Adrian C. Sleigh, G. Richard Olds, Yuesheng Li, Gail M. Williams, and Donald P. McManus. Schistosomiasis. New England Journal of Medicine, 346(SUPPL.103):1212–1220, April 2002. Publisher: World Health Organization ISBN: 0471142735.

17. Yoshiki Aoki, Katsuyuki Sato, N.D. Muhoho, Shin-ichi Noda, and Eisaku Kimura. Cercar-iometry for detection of transmission sites for schistosomiasis. Parasitology International, 52(4):403–408, December 2003.

18. J. H. Ouma, R. F. Sturrock, R. K. Klumpp, and H. C. Kariuki. A comparative evaluation of snail sampling and cereariometry to detect Schistosoma mansoni transmission in a large-scale, longitudinal field-study in Machakos, Kenya. Parasitology, 99(3):349–355, 1989.

19. Sebastian Brachs and Wilfried Haas. Swimming behaviour of Schistosoma mansoni cercariae: responses to irradiance changes and skin attractants. Parasitology Research, 102(4):685–690, March 2008.

20. Simone Haeberlein and Wilfried Haas. Chemical attractants of human skin for swimming Schistosoma mansoni cercariae. Parasitology Research, 102(4):657–662, 2008.

21. C. J. Shiff and Thaddeus K Graczyk. A chemokinetic response in Schistosoma mansoni cercariae. Journal of Parasitology, 80(6):879–883, 1994.

22. Harold L. Asch. Effect of selected chemical agents on longevity and infectivity of Schistosoma mansoni cercariae. Experimental Parasitology, 38(2):208–216, October 1975.

23. Deepak Krishnamurthy, Georgios Katsikis, Arjun Bhargava, and Manu Prakash. Schistosoma mansoni cercariae swim efficiently by exploiting an elastohydrodynamic coupling. Nature Physics, 13(3):266–271, 2017.

24. Kenneth S. Saladin. Schistosoma mansoni: Cercarial Responses to Irradiance Changes. The Journal of Parasitology, 68(1):120, February 1982.

25. Peter J Menck, Jobst Heitzig, Norbert Marwan, and Jürgen Kurths. How basin stability complements the linear-stability paradigm. Nature physics, 9(2):89–92, 2013.

26. SH Lee, RS Chadwick, and L Gary Leal. Motion of a sphere in the presence of a plane interface. part 1. an approximate solution by generalization of the method of lorentz. Journal of Fluid Mechanics, 93(4):705–726, 1979.

27. SH Lee and LG Leal. Motion of a sphere in the presence of a plane interface. part 2. an exact solution in bipolar co-ordinates. Journal of Fluid Mechanics, 98(1):193–224, 1980.

28. Jocelyn Dunstan, Gastón Mino, Eric Clement, and Rodrigo Soto. A two-sphere model for bacteria swimming near solid surfaces. Physics of Fluids, 24(1), 2012.

29. Hoa Nguyen, Amelia Gibbs, Frank Healy, Orrin Shindell, Ricardo Cortez, Kathleen M Brown, Jonathan McCoy, and Bruce Rodenborn. Using theory and experiments of spheres moving near boundaries to optimize the method of images for regularized stokeslets. Physical Review Fluids, 10(3):033101, 2025.

30. Sangtae Kim and Seppo J Karrila. Microhydrodynamics: principles and selected applications. Butterworth-Heinemann, 2013.

31. James W Swan and John F Brady. Simulation of hydrodynamically interacting particles near a no-slip boundary. Physics of Fluids, 19(11), 2007.

32. James W Swan and John F Brady. Particle motion between parallel walls: Hydrodynamics and simulation. Physics of Fluids, 22(10), 2010.

33. Maciej Lisicki, Bogdan Cichocki, and Eligiusz Wajnryb. Near-wall diffusion tensor of an axisymmetric colloidal particle. The Journal of Chemical Physics, 145(3), 2016.

34. Rodrigo Ledesma-Aguilar and Julia M Yeomans. Enhanced motility of a microswimmer in rigid and elastic confinement. Physical review letters, 111(13):138101, 2013.

35. Jörn Dunkel, Victor B Putz, Irwin M Zaid, and Julia M Yeomans. Swimmer-tracer scattering at low reynolds number. Soft Matter, 6(17):4268–4276, 2010.

36. GP Alexander, CM Pooley, and JM Yeomans. Hydrodynamics of linked sphere model swimmers. Journal of Physics: Condensed Matter, 21(20):204108, 2009.

37. Emiliya Passov and Yizhar Or. Dynamics of purcell’s three-link microswimmer with a passive elastic tail. The European Physical Journal E, 35:1–9, 2012.

38. Sankalp Nambiar and John S Wettlaufer. Hydrodynamics of slender swimmers near deformable interfaces. Physical Review Fluids, 7(5):054001, 2022.

39. G R Fulford and JR Blake. On the motion of a slender body near an interface between two immiscible liquids at very low reynolds numbers. Journal of Fluid Mechanics, 127:203–217, 1983.

40. Seung-Man Yang and L Gary Leal. Particle motion in stokes flow near a plane fluid-fluid interface. part 1. slender body in a quiescent fluid. Journal of Fluid Mechanics, 136:393–421, 1983.

41. Lyndon Koens and Thomas D Montenegro-Johnson. Local drag of a slender rod parallel to a plane wall in a viscous fluid. Physical Review Fluids, 6(6):064101, 2021.

42. Jian Teng, Bhargav Rallabandi, Howard A Stone, and Jesse T Ault. Coupling of translation and rotation in the motion of finite-length rods near solid boundaries. Journal of Fluid Mechanics, 938:A30, 2022.

43. Juho S Lintuvuori, Aidan T Brown, Kevin Stratford, and Davide Marenduzzo. Hydrodynamic oscillations and variable swimming speed in squirmers close to repulsive walls. Soft Matter, 12(38):7959–7968, 2016.

44. Kenta Ishimoto and Eamonn A Gaffney. Squirmer dynamics near a boundary. Physical Review E—Statistical, Nonlinear, and Soft Matter Physics, 88(6):062702, 2013.

45. Darren Crowdy. Treadmilling swimmers near a no-slip wall at low reynolds number. International Journal of Non-Linear Mechanics, 46(4):577–585, 2011.

46. Allison P Berke, Linda Turner, Howard C Berg, and Eric Lauga. Hydrodynamic attraction of swimming microorganisms by surfaces. Physical Review Letters, 101(3):038102, 2008.

47. Eric Lauga, Willow R DiLuzio, George M Whitesides, and Howard A Stone. Swimming in circles: motion of bacteria near solid boundaries. Biophysical journal, 90(2):400–412, 2006.

48. David F Katz. On the propulsion of micro-organisms near solid boundaries. Journal of Fluid Mechanics, 64(1):33–49, 1974.

49. Jens Elgeti and Gerhard Gompper. Self-propelled rods near surfaces. Europhysics Letters, 85(3):38002, 2009.

50. Darren G Crowdy and Yizhar Or. Two-dimensional point singularity model of a low-reynolds-number swimmer near a wall. Physical Review E—Statistical, Nonlinear, and Soft Matter Physics, 81(3):036313, 2010.

51. Yizhar Or and Richard M Murray. Dynamics and stability of a class of low reynolds number swimmers near a wall. Physical Review E—Statistical, Nonlinear, and Soft Matter Physics, 79(4):045302, 2009.

52. Yizhar Or. Dynamics and stability of purcell’s three-link microswimmer near a wall. Physical Review E—Statistical, Nonlinear, and Soft Matter Physics, 82(6):065302, 2010.

53. Renaud Trouilloud, Tony S Yu, AE Hosoi, and Eric Lauga. Soft swimming: exploiting deformable interfaces for low reynolds number locomotion. Physical review letters, 101(4):048102, 2008.

54. Darren Crowdy, Sungyon Lee, Ophir Samson, Eric Lauga, and AE Hosoi. A two-dimensional model of low-reynolds number swimming beneath a free surface. Journal of Fluid Mechanics, 681:24–47, 2011.

55. Vaseem A Shaik and Arezoo M Ardekani. Motion of a model swimmer near a weakly deforming interface. Journal of Fluid Mechanics, 824:42–73, 2017.

56. Roberto Di Leonardo, Dario Dell’Arciprete, Luca Angelani, and Valerio Iebba. Swimming with an image. Physical review letters, 106(3):038101, 2011.

57. Sungyon Lee, John WM Bush, AE Hosoi, and Eric Lauga. Crawling beneath the free surface: Water snail locomotion. Physics of Fluids, 20(8), 2008.

58. Saverio E Spagnolie and Eric Lauga. Hydrodynamics of self-propulsion near a boundary: predictions and accuracy of far-field approximations. Journal of Fluid Mechanics, 700:105–147, 2012.

59. S Wang and AM Ardekani. Swimming of a model ciliate near an air-liquid interface. Physical Review E—Statistical, Nonlinear, and Soft Matter Physics, 87(6):063010, 2013.

60. Diego Lopez and Eric Lauga. Dynamics of swimming bacteria at complex interfaces. Physics of Fluids, 26(7), 2014.

61. Zhengbin Zhang, Liansheng Liu, Chunying Liu, and Weijun Cai. Studies on the sea surface microlayer. Journal of Colloid and Interface Science, 264(1):148–159, August 2003.

62. J. Ruth Lawson, R. A. Avilson, and J. Ruth Lawson. The survival of the cercariae of Schistosoma mansoni in relation to water temperature and glycogen utilization. Parasitology, 81(2):337–348, 1980.

63. Shiqiang Wang, Yongqi Shi, and Li Wen. Physical Intelligence in Biomechanics. IOP Conference Series: Materials Science and Engineering, 1261(1):012012, October 2022.

64. F Haas, S Gorb, and R.J Wootton. Elastic joints in dermapteran hind wings: materials and wing folding. Arthropod Structure & Development, 29(2):137–146, April 2000.

65. Danli Luo, Aditi Maheshwari, Andreea Danielescu, Jiaji Li, Yue Yang, Ye Tao, Lingyun Sun, Dinesh K. Patel, Guanyun Wang, Shu Yang, Teng Zhang, and Lining Yao. Autonomous self-burying seed carriers for aerial seeding. Nature, 614(7948):463–470, February 2023.

66. D. N. Beal, F. S. Hover, M. S. Triantafyllou, J. C. Liao, and G. V. Lauder. Passive propulsion in vortex wakes. Journal of Fluid Mechanics, 549(-1):385, February 2006.

67. Brooke J. Pauken, Andrew J. Chamberlin, Chelsea L. Wood, Oliver B. Fringer, and Giulio A. De Leo. Modeling of wind-driven circulation of schistosome larvae in a vegetated side pond. Environmental Fluid Mechanics, 25(2):13, April 2025.

68. Jason P. Sckrabulis, Alan R. Flory, and Thomas R. Raffel. Direct onshore wind predicts daily swimmer’s itch (avian schistosome) incidence at a Michigan beach. Parasitology, 147(4):431– 440, April 2020.

69. Jingran Qiu, Cristian Marchioli, and Lihao Zhao. A review on gyrotactic swimmers in turbulent flows. Acta Mechanica Sinica, 38(8):722323, August 2022.

70. William M. Durham, Eric Climent, Michael Barry, Filippo De Lillo, Guido Boffetta, Massimo Cencini, and Roman Stocker. Turbulence drives microscale patches of motile phytoplankton. Nature Communications, 4(1):2148, July 2013.

71. William M. Durham, John O. Kessler, and Roman Stocker. Disruption of Vertical Motility by Shear Triggers Formation of Thin Phytoplankton Layers. Science, 323(5917):1067–1070, February 2009.

72. Fabian Reitzug, Narcis B. Kabatereine, Anatol M. Byaruhanga, Fred Besigye, Betty Nabatte, and Goylette F. Chami. Current water contact and Schistosoma mansoni infection have distinct determinants: a data-driven population-based study in rural Uganda. Nature Communications, 15(1):9530, November 2024. Publisher: Nature Publishing Group.

73. Jan Pieter R. Koopman, Emma L. Houlder, Jacqueline J. Janse, Miriam Casacuberta-Partal, Olivia A.C. Lamers, Jeroen C. Sijtsma, Claudia de Dood, Stan T. Hilt, Arifa Ozir-Fazalalikhan, Vincent P. Kuiper, Geert V.T. Roozen, Laura M. de Bes-Roeleveld, Yvonne C.M. Kruize, Linda J. Wammes, Hermelijn H. Smits, Lisette van Lieshout, Govert J. van Dam, Inge M. van Amerongen-Westra, Pauline Meij, Paul L.A.M. Corstjens, Simon P. Jochems, Angela van Diepen, Maria Yazdanbakhsh, Cornelis H. Hokke, and Meta Roestenberg. Safety and infectivity of female cercariae in Schistosoma-naïve, healthy participants: a controlled human Schistosoma mansoni infection study. eBioMedicine, 97:104832, October 2023.

74. A. C. Fusco, L. Cassioppi, B. Salafsky, and T. Shibuya. Penetration of Schistosoma mansoni cercariae into a living skin equivalent. The Journal of Parasitology, 79(3):444–448, June 1993.

75. Allen W. Cheever. A Quantitative Post-Mortem Study of Schistosomiasis Mansoni in Man. The American Journal of Tropical Medicine and Hygiene, 17(1):38–64, January 1968. Publisher: The American Society of Tropical Medicine and Hygiene Section: The American Journal of Tropical Medicine and Hygiene.

76. A. J. C. Fulford, A. E. Butterworth, J. H. Ouma, and R. F. Sturrock. A statistical approach to schistosome population dynamics and estimation of the life-span of Schistosoma mansoni in man. Parasitology, 110(3):307–316, April 1995.

77. Pmz Coelho and Rl Caldeira. Critical analysis of molluscicide application in schistosomiasis control programs in Brazil. Infectious Diseases of Poverty, 5(1):57, December 2016.

78. F. A. Lewis, M. A. Stirewalt, C. P. Souza, and G. Gazzinelli. Large-scale laboratory maintenance of Schistosoma mansoni, with observations on three schistosome/snail host combinations. The Journal of Parasitology, 72(6):813–829, December 1986.

79. Matthew S. Tucker, Laksiri B. Karunaratne, Fred A. Lewis, Tori C. Freitas, and Yung-San Liang. Schistosomiasis. Current Protocols in Immunology, 103:19.1.1–19.1.58, November 2013.

80. Dmitry Ershov, Minh-Son Phan, Joanna W. Pylvänäinen, Stéphane U. Rigaud, Laure Le Blanc, Arthur Charles-Orszag, James R. W. Conway, Romain F. Laine, Nathan H. Roy, Daria Bonazzi, Guillaume Duménil, Guillaume Jacquemet, and Jean-Yves Tinevez. Bringing TrackMate into the era of machine-learning and deep-learning, September 2021. Pages: 2021.09.03.458852 Section: New Results.

81. Johannes Schindelin, Ignacio Arganda-Carreras, Erwin Frise, Verena Kaynig, Mark Longair, Tobias Pietzsch, Stephan Preibisch, Curtis Rueden, Stephan Saalfeld, Benjamin Schmid, Jean-Yves Tinevez, Daniel James White, Volker Hartenstein, Kevin Eliceiri, Pavel Tomancak, and Albert Cardona. Fiji: an open-source platform for biological-image analysis. Nature Methods, 9(7):676–682, July 2012. arXiv: 1081-8693 ISBN: 1548-7105 (Electronic)\r1548-7091 (Linking).

